# GRK2/3/5/6 knockout: The impact of individual GRKs on arrestin-binding and GPCR regulation

**DOI:** 10.1101/2021.02.12.430971

**Authors:** J. Drube, R.S. Haider, E.S.F. Matthees, M. Reichel, J. Zeiner, S. Fritzwanker, C. Ziegler, S. Barz, L. Klement, A. Kliewer, E. Miess, E. Kostenis, S. Schulz, C. Hoffmann

**Author notes:** To whom correspondence should be addressed: Prof. Dr. Carsten Hoffmann, Institut für Molekulare Zellbiologie, CMB – Center for Molecular Biomedicine, Universitätsklinikum Jena, Friedrich-Schiller-Universität Jena, Hans-Knöll-Straße 2, D-07745 Jena, Germany. contributed equally (J. Drube, R.S. Haider).

## Abstract

G protein-coupled receptors (GPCRs) comprise the largest family of transmembrane receptors and represent major drug targets. Upon ligand stimulation, GPCRs activate G proteins and undergo a complex regulation by interaction with GPCR kinases (GRKs) and formation of receptor–arrestin complexes. For many GPCRs, this mechanism triggers receptor desensitisation, internalisation, and possibly a second intracellular signalling wave. Here we created eleven different HEK293 knockout cell clones for GRK2, 3, 5, and 6 individually and in combination. These include four single, two double, four triple, and the quadruple GRK knockout. The statistical evaluation of β-arrestin1/2 interactions for twelve different receptors grouped the tested GPCRs into two main subsets: those for which β-arrestin interaction was mediated by either GRK2, 3, 5, or 6 and those that are mediated by GRK2 or 3 only. Interestingly, the overexpression of specific GRKs was found to induce a robust, ligand-independent β-arrestin interaction with the V2R and AT1R. Finally, using GRK knockout cells, PKC inhibitors, and β-arrestin mutants, we present evidence for differential AT1R–β-arrestin2 complex configurations mediated by selective engagement of PKC, GRK2, or GRK6. We anticipate our novel GRK-knockout platform to facilitate the elucidation of previously unappreciated details of GRK-specific GPCR regulation and β-arrestin complex formation.

## Introduction

G protein-coupled receptors (GPCRs) constitute the largest family of membrane receptors in human physiology comprising more than 800 identified members. GPCRs regulate multitudes of physio- and pathophysiological processes and are well established targets for pharmacological intervention. In fact, a recent review listed 134 GPCRs and about 50 additional GPCR-signalling related proteins which are directly targeted by FDA-approved drugs^1^.

The diverse stimuli recognized by GPCRs induce conformational changes within the receptor, which activate distinct signalling pathways^2^. As opposed to the large number of GPCRs, the intracellular signalling molecules are less diverse. Besides G proteins, GPCR kinases (GRKs) and arrestins are the most immediate GPCR-interacting molecules^3,4^. In the non-visual system only four GRKs and two arrestin isoforms are hypothesised to regulate hundreds of GPCRs^5^. To accommodate for this apparent imbalance, the phosphorylation barcode hypothesis for receptor–arrestin interactions was developed^6–8^. Since GRK-induced GPCR phosphorylation is the basis for high-affinity β-arrestin binding, we anticipate that individual GRK isoforms shape the GPCR signalling response in a cell and tissue-specific manner.

The relative selectivity of ligands to favour a certain pathway at the expense of others was termed functional selectivity or biased agonism^5,9,10^. The recognition that either G protein- or arrestin-supported pathways^11^ can contribute to pathophysiological conditions or drug-associated side effects^9,12^ triggered an intense search for biased ligands. GRKs act as essential mediators and define β-arrestin functions via ligand-specific GPCR phosphorylation or preferential coupling to certain active receptor states. Structural biology greatly contributed to our understanding of receptor conformational changes which lead to the interaction with either G proteins or arrestins. A multistep interaction mechanism for arrestins involving receptor phosphorylation^13^ and the arrestin finger loop region^14,15^ has been revealed. However, little is currently known about the impact of individual GRKs on arrestin binding.

The ubiquitous expression of GRK2, 3, 5, and 6 obscures the elucidation of the roles of individual GRKs on receptor phosphorylation. Until now, siRNA/shRNA^16–18^, or CRISPR/Cas 9 approaches targeting only a certain subset of relevant GRKs^19^, and the utilisation of GRK inhibitors were the only strategies used to study their impact on living cell function. Yet, in combination with phosphosite-specific antibodies^20,21^ or mass-spectrometry^22^, contributions of individual GRKs to the phosphorylation of certain receptors were elucidated to some degree^7^.

Nevertheless, the remaining expression of the targeted GRK(s) in knockdown approaches, or potential off-target effects of pharmacological intervention preclude the unambiguous interpretation of obtained results. Thus, a comprehensive elucidation of single GRK contributions to the arrestin-dependent regulation of GPCR signalling, internalisation, and trafficking remains elusive.

In this study, we present a cellular platform to investigate the individual roles of GRK2, 3, 5, and 6 in these processes. We have created a panel of eleven combinatorial HEK293 GRK knockout clones, which enable us to analyse the GRK contributions to GPCR phosphorylation, recruitment of β-arrestin1 and 2, as well as receptor internalisation in unprecedented detail.

### GRK knockout cells: a viable cellular platform to assess individual GRK contributions

Utilising the CRISPR/Cas9 technology, we engineered HEK293 single-cell clones with knockouts (KO) of GRKs. We created single KOs of GRK2 (ΔGRK2), GRK3 (ΔGRK3), GRK5 (ΔGRK5), GRK6 (ΔGRK6), two double KOs ΔGRK2/3 and ΔGRK5/6, and a quadruple KO of GRK2, 3, 5, and 6 (ΔQ-GRK). Additionally, we established four triple KO cell clones (ΔGRK3/5/6, ΔGRK2/5/6, ΔGRK2/3/6, ΔGRK2/3/5) with the endogenous expression of one remaining GRK. The KOs were confirmed by western blot analysis **(Figure 1a)** and as recommended^23^, further confirmed by functional studies in this manuscript. Morphology as revealed by phase-contrast microscopy of cultured cells and cell growth **(Supplementary Figure 1a, b)** were only mildly affected in some of the clones. Expression levels of the untargeted GRKs in the obtained cell clones remained virtually unchanged compared to Control **(Supplementary Figure 1c)**.

**Figure 1:**
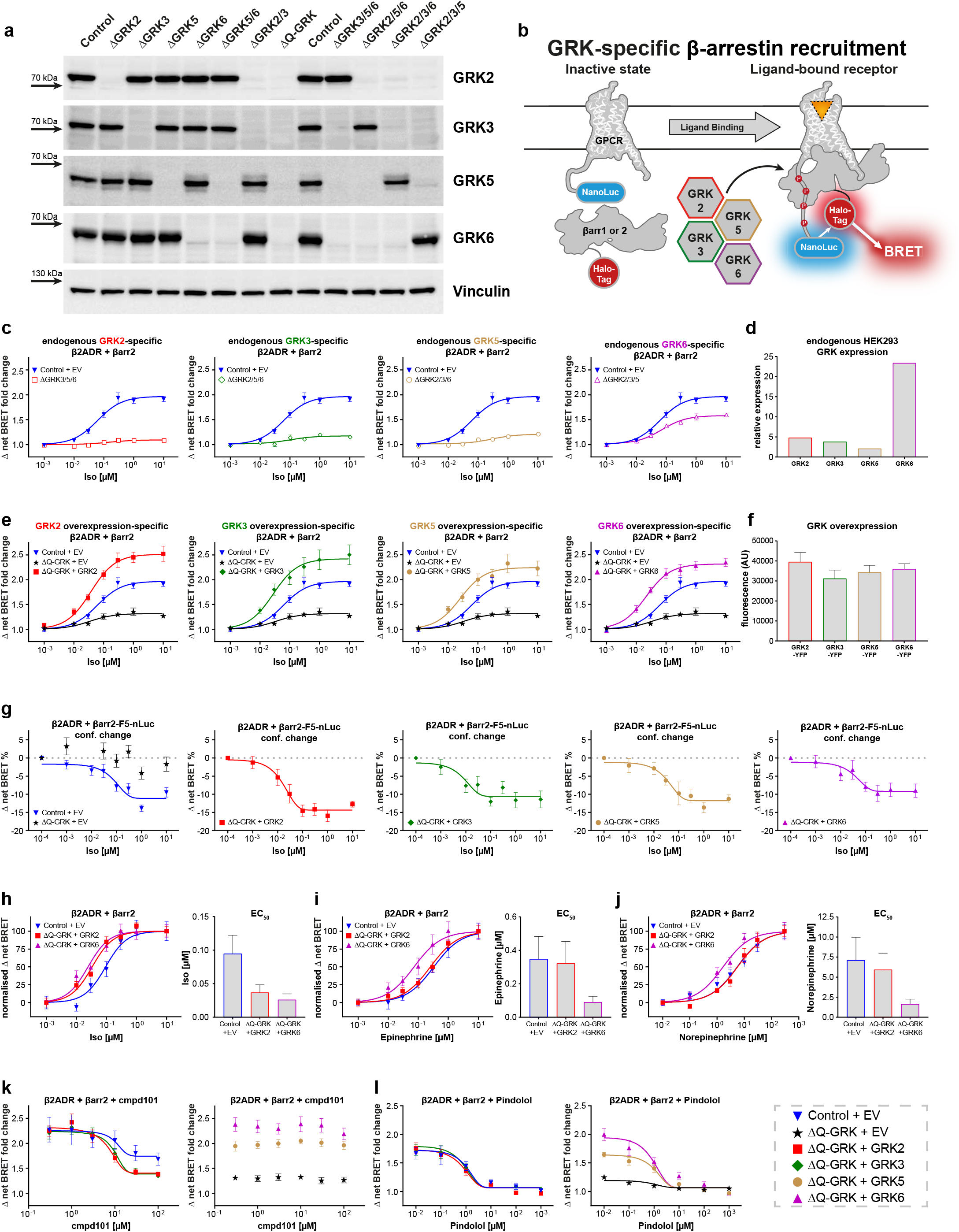
GRK knockout cells: a viable cellular platform to assess the GRK-specific formation of β2ADR–β-arrestin complexes. **a** Single (ΔGRK2, 3, 5, 6), double (ΔGRK2/3 or 5/6), triple (ΔGRK3/5/6, 2/5/6, 2/3/6, 2/3/5), and quadruple (ΔQ-GRK) cells were generated using the CRISPR/Cas9 technology and established as single cell clones. The absence of GRK2, 3, 5, or 6 was confirmed by western blot analysis (for quantification, see **Supplementary Figure 1c**). **b** Schematic depiction of the performed NanoBRET β-arrestin (βarr) recruitment assay and colour-coding for GRK-specific conditions used throughout the manuscript. The Halo-Tag-βarr fusion protein is recruited to a NanoLuciferase (NanoLuc)-tagged GPCR upon agonist activation and subsequent receptor phosphorylation. The resulting change in proximity of the Halo-Tag and the NanoLuc increases measured BRET ratios, enabling the agonist concentration-dependent analysis of βarr recruitment. **c** Halo-Tag-βarr2 recruitment to the β2ADR-NanoLuc upon stimulation with isoprenaline (Iso) in cells expressing all endogenous GRKs (Control + empty vector (EV)) or only one remaining endogenous GRK (triple knockout cells ΔGRK3/5/6, ΔGRK2/5/6, ΔGRK2/3/6, ΔGRK2/3/5). **d** The normalised endogenous expression levels of all four GRKs in HEK293 cells^26^, as compiled by the human protein atlas (http://proteinatlas.org/humanproteome/cell). **e** βarr2 recruitment to the β2ADR in quadruple GRK knockout cells (ΔQ-GRK), overexpressing a single GRK (ΔQ-GRK + GRK) or expressing all endogenous GRKs (Control + EV). BRET data in **(c)** and **(e)** are presented as Δ net BRET fold change, mean of three independent experiments ± SEM. For better comparison, the Control and ΔQ-GRK curves are shown multiple times. **f** GRK-YFP fusion proteins were transfected in ΔQ-GRK and YFP fluorescence was measured to confirm similar expression levels of all transfected GRKs. Corresponding experiments, confirming the catalytic activity of GRK-YFP fusion constructs are shown in **Supplementary Figure 2a**. Measured fluorescence is depicted as mean + SD of four independent transfections, recorded on two measuring days as arbitrary units (AU). **g** Analysis of βarr conformational changes. ΔQ-GRK or Control cells were transfected with an untagged β2ADR expression construct and the βarr2-F5-NanoLuc conformational change biosensor (more details on the novel BRET sensor will be published elsewhere), in absence or presence of GRKs as noted and stimulated with Iso. Conformational change data are shown as Δ net BRET change in percent, mean of three independent repetitions ± SEM. **h-j** The recruitment of βarr2 to the β2ADR following stimulation with Iso **(h)**, Epinephrine **(i)**, or Norepinephrine **(j)** in presence of all endogenous GRKs or individually overexpressed GRK2 or GRK6 in ΔQ-GRK as indicated. Data are depicted as mean of three independent experiments ± SEM and normalised to individual maxima. The bar graphs represent the EC50 + SEM of the corresponding concentration-response curves. **k, l** Utilisation of the βarr recruitment assay presented in **(e)** for specificity determination of the GRK inhibitor cmpd101 in living cells. ΔQ-GRK or Control cells were transfected with β2ADR-NanoLuc, Halo-Tag-βarr2, and either GRK2, 3, 5, 6, or EV as noted. The cells were incubated with different concentrations of cmpd101 **(k)** or the β2ADR antagonist pindolol **(l)** for 10 minutes prior to stimulation with 1 μM Iso. The recruitment-induced BRET changes were measured and calculated as Δ net BRET change in percent, represented as the mean ± SEM of four independent experiments.

In order to test the effect of our ΔGRK-clones and confirm the unaltered kinase activity of GRKs untargeted in a specific KO cell line, we revisited and analysed agonist promoted μ-opioid receptor (MOP) phosphorylation. This receptor system was deliberately chosen, as GRK contributions were already extensively studied^19,24^ and the abundance of available phosphosite-specific antibodies allowed for a complete elucidation of D-Ala^2^, N-MePhe^4^, Gly-ol]-enkephalin (DAMGO) induced receptor phosphorylation **(Supplementary Figure 1d, e)**. We successfully identified T^376^ as a specific target of GRK2 and 3. In line with our previous findings^24^, T^370^, S^375^, and T^379^ seem to be phosphorylated by all four GRKs to different extents. In ΔQ-GRK, the phosphorylation of these sites was completely abolished, confirming them as GRK target sites. In contrast, S^363^, a known PKC phosphorylation site^24,25^ showed a strong phosphorylation signal in ΔQ-GRK cells **(Supplementary Figure 1d, e)**. This analysis confirms the general functionality of our clones for phosphorylation studies and underlines the unaltered activity of kinases not targeted in our KO approach.

Further we investigated the internalisation of the MOP in Control-, ΔGRK2/3, ΔGRK5/6, and ΔQ-GRK stably expressing the MOP by confocal microscopy and surface enzyme-linked immunosorbent assay (ELISA) **(Supplementary Figure 1f, g)**. After stimulation, the MOP was internalised in Control and ΔGRK5/6, but remained at the cell surface in ΔGRK2/3 and ΔQ-GRK. Our findings confirm that all four analysed GRKs are able to act on the MOP, but exclusively GRK2 and 3 are able to phosphorylate T^376^ and further drive internalisation of the receptor.

### GRK2, 3, 5, and 6 are able to individually induce the formation of β2ADR–β-arrestin complexes

As the availability of tools for the analysis of site-specific receptor phosphorylation is limited across the GPCR superfamily, we utilised the universal GPCR adaptor proteins β-arrestin1 and 2 to analyse the contributions of individual GRKs to receptor regulation. The schematic in **Figure 1b** depicts the established bioluminescence resonance energy transfer (BRET)-based *in cellulo* β-arrestin recruitment assay, allowing us to reveal functional, GRK-specific GPCR phosphorylation.

First, we studied the GRK-specific interactions between the β2 adrenergic receptor (β2ADR) and β-arrestin2 utilising the endogenous expression of GRKs in various ΔGRK cells. Under the endogenous expression of all four GRKs (Control + EV), β-arrestin2 showed a clear isoprenaline-(Iso) induced recruitment to the β2ADR **(Figure 1c)**. In comparison, the measured β-arrestin recruitment was drastically reduced when recorded in triple GRK KO cell lines, only featuring the endogenous expression of one individual GRK (**Figure 1c**, ΔGRK3/5/6, ΔGRK2/5/6, ΔGRK2/3/6, ΔGRK2/3/5). Surprisingly, the endogenous expression of either GRK2, 3, or 5 induced only minimal BRET changes. Nevertheless, we were able to detect a ligand-dependent increase in β-arrestin recruitment. In this study, data which could be described by a curve-fit will be further interpreted as functional recruitment.

The highest amount of β-arrestin2 recruitment was found in ΔGRK2/3/5 cells, specifically induced by the endogenous expression of GRK6. This finding correlates with the relative abundance of GRKs in HEK293 cells^26^, as compiled by the human protein atlas (http://proteinatlas.org/humanproteome/cell) **(Figure 1d)**, with GRK6 being approximately 5-fold more expressed than the other assessed isoforms. Thus, we were able to identify GRK6 as the main mediator of Iso-promoted β-arrestin recruitment to the β2ADR, when measured under endogenous expression of GRKs in HEK293 cells.

Since these findings specifically reflected on the expression levels of GRKs, we analysed the molecular capability of individually over-expressed GRKs to induce β2ADR–β-arrestin2 complex formation via re-introduction of GRK2, 3, 5, or 6 into ΔQ-GRK **(Figure 1e)**. The relative expression of transfected GRKs was fluorometrically assessed in **Figure 1f**. Via introduction of a C-terminal YFP fusion into the identical vector backbone and subsequent equimolar transfection of GRK-YFP constructs, we confirmed similar expression levels of the transfected kinases. To allow for this comparison, the GRK-YFP fusion proteins were characterised with at least the same catalytic activity to mediate GPCR–β-arrestin interactions as their untagged counterparts, used in all other experiments **(Supplementary Figure 2a)**. Using this controlled overexpression of individual GRKs in ΔQ-GRK, all four kinases showed a similar effect on β2ADR regulation: any individual GRK isoform can enhance the β2ADR–β-arrestin recruitment to higher levels than induced by the combined endogenous expression of GRKs in Control + EV **(Figure 1e)**. Interestingly, we still encountered measurable β-arrestin2 recruitment in the absence of GRKs (ΔQ-GRK + EV). This could be explained by the inherit affinity of β-arrestin2 towards ligand-activated, yet unphosphorylated GPCRs, or the action of other kinases.

These findings clarify that all four tested GRKs are able to individually mediate high-affinity β-arrestin2 binding to the β2ADR and their relative tissue expression ultimately defines their specific contributions to this process.

Since all GRKs have been shown to induce similar levels of β-arrestin recruitment, we investigated whether subtype-specific phosphorylation of the β2ADR might still have a pronounced effect on the conformational changes that occur during arrestin activation. Here, the single overexpression of all GRK isoforms alongside the untagged β2ADR induced comparable β-arrestin2 conformational changes when measured with the β-arrestin2-FlAsH5-NanoLuc BRET biosensor, originally described as a FRET sensor in Nuber *et al.^27^* **(Figure 1g,** more details on the novel BRET sensor will be published elsewhere).

We further utilised the β2ADR, as a model receptor regulated by all four tested GRK isoforms, to test the effect of endogenous ligands and pharmacological inhibition on GRK-specific β-arrestin-coupling processes. The application of the endogenous ligands epinephrine and norepinephrine resulted in overall lower GRK-specific β-arrestin2 recruitment to the β2ADR as compared to Iso **(Supplementary Figure 2b, c** compare with **Figure 1e)**. Although the relative efficacies of individual GRKs to mediate epinephrine- and norepinephrine-induced β-arrestin2 binding were unchanged in comparison to Iso (ΔQ-GRK + GRK > Control + EV > ΔQ-GRK + EV), we observed a left shift of the measured concentration-response curves specifically for GRK6 **(Figure 1h, i, j)**. This apparent increase in potency was observed for both endogenous ligands, although it was shown that epinephrine acts as a full agonist, whereas norepinephrine only partially activates the β2ADR^28^.

Furthermore, utilising the interaction between the β2ADR and β-arrestin2, we established a cell-based GRK-inhibitor screening platform. We were able to record the concentration-dependent inhibition of β-arrestin2 recruitment to the receptor by cmpd101 (a known GRK2 family inhibitor) only in ΔQ-GRK overexpressing GRK2 or 3 as well as in Control **(Figure 1k)**. Thus, we demonstrate cmpd101 selectivity in living cells by the lack of inhibition when overexpressing GRK5 or 6. When performing the analogous experiment using pindolol as a potent antagonist of the β2ADR, we recorded an inhibition of β-arrestin2 recruitment regardless of GRK (over-) expression **(Figure 1l)**.

### ΔQ-GRK cells reveal GRK-specificity of β-arrestin1 and 2 recruitment to different GPCRs

To compare the individual capabilities of GRK2, 3, 5 and 6 to facilitate β-arrestin recruitment across multiple receptors, we analysed a total of twelve different GPCRs. The panel was chosen deliberately to feature receptors with divergent lengths of intracellular domains to achieve a comprehensive overview of GRK-selective β-arrestin1 and 2 recruitment: angiotensin 1 receptor (AT1R), β2ADR, β2ADR with an exchanged C-terminus of the vasopressin 2 receptor (β2V2), complement 5a receptor 1 (C5aR1), muscarinic acetylcholine receptors (M1R, M2R, M3R, M4R, M5R), MOP, parathyroid hormone 1 receptor (PTH1R), and vasopressin 2 receptor (V2R). **(Figure 2a-d; Supplementary Figures 3, 4; Supplementary Table 1)**.

**Figure 2:**
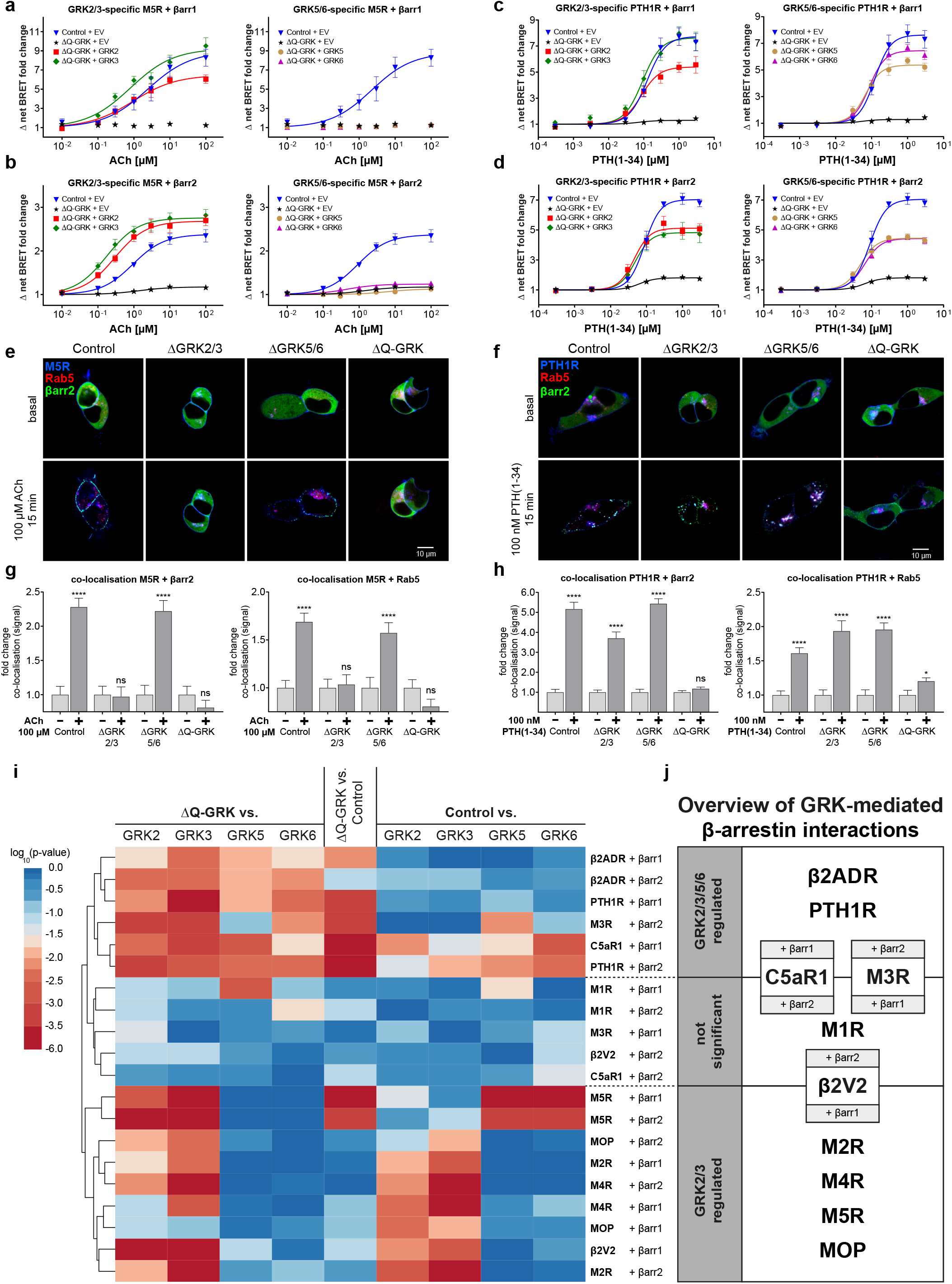
ΔQ-GRK cells reveal GRK-specificity of β-arrestin1 and 2 recruitment to different GPCRs and allow assessment of GRK-dependent GPCR internalisation and β-arrestin2 translocation. **a-d** GRK-specific β-arrestin (βarr)1 **(a, c)** or βarr2 **(b, d)** recruitment to the M5R upon acetylcholine (ACh) stimulation **(a, b)** or the PTH1R upon parathyroid-hormone 1-34 (PTH(1-34)) stimulation **(c, d)**. Shown are concentration-response curves depicted as Δ net BRET fold change. The panels display the recruitment in presence of either GRK2 or 3 or GRK5 or 6. For better comparison, the Control and ΔQ-GRK curves are shown multiple times. **e, f** Control, ΔGRK2/3, ΔGRK5/6, and ΔQ-GRK cells were transfected with either M5R-CFP or PTH1R-CFP (blue), the early endosome marker Rab5-mCherry (red), and βarr2-YFP (green) expression constructs. The cells were grown on cover slips and subjected to confocal live-cell microscopy. Shown are representative images, taken before and after 15 minutes of stimulation with either 100 μM ACh or 100 nM PTH(1-34), respectively. The normalised co-localisation of M5R **(g)** or PTH1R **(h)** with βarr2 or Rab5 was quantified using Squassh and SquasshAnalyst, with at least 30 images per condition, representing at least three independent transfections and experimental days. Data are presented as mean fold change in co-localisation signal + SEM. Statistical analysis was performed using a two-way mixed model ANOVA followed by a paired t-test (*p < 0.05; ** p < 0.01; *** p < 0.001; **** p < 0.0001; ns (not significant)). **i** Clustering heatmap representing the statistical multiple comparison of βarr recruitment data for ten different GPCRs. Conditions with overexpressed GRKs were tested against ΔQ-GRK + empty vector (EV) or Control + EV, as indicated. Additionally ΔQ-GRK + EV was compared to Control + EV. BRET fold changes at saturating ligand concentrations of at least three independent experiments were compared using ANOVA and Bonferroni’s test **(Supplementary Table 2,** data derived from **Supplementary Figure 3)**. Transformed unadjusted p values are plotted. GPCR–βarr pairs are clustered according to Canberra distance. **j** Overview of clustering from **(i)** in GPCR–βarr pairs regulated by any tested GRK (GRK2/3/5/6 regulated), by GRK2 or 3 only (GRK2/3 regulated) and a third group, which is comprised of GPCR–βarr pairs that do not consistently show significant differences between the tested conditions.

As representative examples of our findings, the GRK-selective β-arrestin1 and 2 recruitment to the M5R and PTH1R are depicted in **Figure 2a, b** and **Figure 2c, d**, respectively. Both receptors were able to induce robust, agonist-dependent β-arrestin1 and 2 recruitment in Control+EV, which was significantly reduced in ΔQ-GRK+EV. In case of the M5R, β-arrestin1 recruitment was completely abolished in ΔQ-GRK+EV. Still, a major difference in GRK-selectivity of the two receptors was found using this approach: the individual overexpression of GRK2, 3, 5, and 6 significantly increased β-arrestin recruitment to the PTH1R in ΔQ-GRK, whereas GRK5 and 6 were unable to facilitate M5R–β-arrestin complex formation. These findings were additionally confirmed in triple GRK KO cell lines and the ΔGRK2/3 and ΔGRK5/6 family KOs **(Supplementary Figure 5a, b)**. Interestingly, the endogenous expression of GRK2 and 3 in ΔGRK3/5/6 and ΔGRK2/5/6 was sufficient to increase the measured β-arrestin2 recruitment in comparison to ΔQ-GRK for both receptors. This finding essentially confirms the functionality of these two triple GRK KO cell lines and suggests that the β2ADR requires higher amounts of GRK2 or 3 in order to be efficiently regulated in comparison to the M5R and PTH1R. As indicated by the experiments shown in **Figure 2b**, ΔGRK cell lines only featuring the expression of GRK5 and/or 6 did not increase the β-arrestin2 recruitment to the M5R as in comparison to ΔQ-GRK.

Further, we employed confocal live-cell microscopy to assess the dependency of PTH1R and M5R internalisation on endogenous GRK levels in Control, ΔGRK2/3, and ΔGRK5/6 as well as in ΔQ-GRK. Under basal conditions, β-arrestin2 is located in the cytosol, M5R and PTH1R in the cell membrane, and Rab5 (early endosome marker) in endosomes **(Figure 2e, f basal)**. As expected, the M5R was not able to induce β-arrestin2 translocation in the absence of GRK2 and 3 (ΔGRK2/3 and ΔQ-GRK) **(Figure 2e)**.

The quantification of co-localisation between the M5R and β-arrestin2 **(Figure 2g)** confirms our findings of **Figure 2b** and **Supplementary Figure 5a**. Analysis of M5R co-localisation with Rab5 (as a surrogate measurement for receptor internalisation and initial trafficking) reveals that this interaction translates to functional receptor internalisation only in the presence of GRK2 and 3 **(Figure 2g)**. For the PTH1R, we were able to detect ligand-induced co-localisation with β-arrestin2 or Rab5 in all conditions expressing GRKs **(Figure 2h)**. Interestingly, the agonist-stimulated PTH1R was still able to induce a slight membrane translocation of β-arrestin2 in ΔQ-GRK, confirming the GRK-independent affinity of β-arrestin2 towards the ligand-activated receptor **(Figure 2d)**. The results obtained for the PTH1R using endogenous GRK expression were verified in a reciprocal experiment overexpressing single GRKs in ΔQ-GRK for β-arrestin1 and 2 **(Supplementary Figure 5c-f)**.

These apparent differences in GRK-specific β-arrestin recruitment, as exemplified by the M5R and the PTH1R, were encountered multiple times during our analysis across twelve different GPCRs **(Supplementary Figure 3)**. Via statistical multiple comparison of BRET fold changes at saturating ligand concentrations for each of the tested conditions (Control + EV, ΔQ-GRK + EV, ΔQ-GRK + GRK2, ΔQ-GRK + GRK3, ΔQ-GRK + GRK5, ΔQ-GRK + GRK6), we were able to cluster the respective GPCR–β-arrestin pairs into groups, depending on the found GRK-selectivity **(Figure 2i, j; Supplementary Table 2)**. Here we identified two main subsets of GPCRs: receptors for which β-arrestin interaction is mediated by overexpression of i) any GRK (β2ADR, PTH1R, C5aR1+β-arrestin1 and M3R+β-arrestin2) or ii) GRK2 or 3 only (M2R, M4R, M5R, MOP, and β2V2+β-arrestin1). Within our tested GPCRs, we did not observe β-arrestin interaction mediated exclusively by GRK5 or 6. Interestingly, some receptor–β-arrestin pairs did not consistently show significant differences between the tested conditions and hence could not be definitively assigned to one of the two groups (M1R, C5aR1+β-arrestin2, M3R+β-arrestin1, and β2V2+β-arrestin2). In case of C5aR1 + β-arrestin2 **(Supplementary Figure 3d)** this behaviour is explained by exceptionally high β-arrestin2 recruitment in the absence of GRKs.

Interestingly, two distinct GPCRs, namely the AT1R and V2R, evaded the statistical grouping process. Both receptors exhibited conditions of diminished, agonist-dependent β-arrestin recruitment in the presence of certain overexpressed GRKs as in comparison to ΔQ-GRK **(Supplementary Figure 3g, m)**. As this finding was highly unexpected, we further focussed on the elucidation of these GRK-dependent processes.

### GRK2, 3, 5, or 6 can individually mediate ligand-independent β-arrestin1 and 2 interactions with the V2R

Besides previous intensive studies^29,30^, the V2R and AT1R presented themselves as distinct cases in our detailed, GRK subtype-specific β-arrestin recruitment assay **(Figure 3, Supplementary Figure 3m)**. In the presence of all four endogenously expressed GRKs, agonist stimulation induced a clear recruitment of β-arrestin1 and 2 to the V2R **(Figure 3a,b; Supplementary Figure 6a, b)**. In ΔQ-GRK the recruitment of β-arrestins was reduced, as expected. Surprisingly, the individual overexpression of GRK2, 3, 5, or 6 did not further increase the concentration-dependent, dynamic change of interaction. When comparing the respective BRET ratios measured before and after stimulation at ligand saturation **(Figure 3a, b; Supplementary Figure 6a, b)**, we found that already the basal BRET ratios were remarkably increased in presence of overexpressed GRKs. This was unlike any receptor presented in **Figure 2i (Supplementary Figure 3, 4)**.

**Figure 3:**
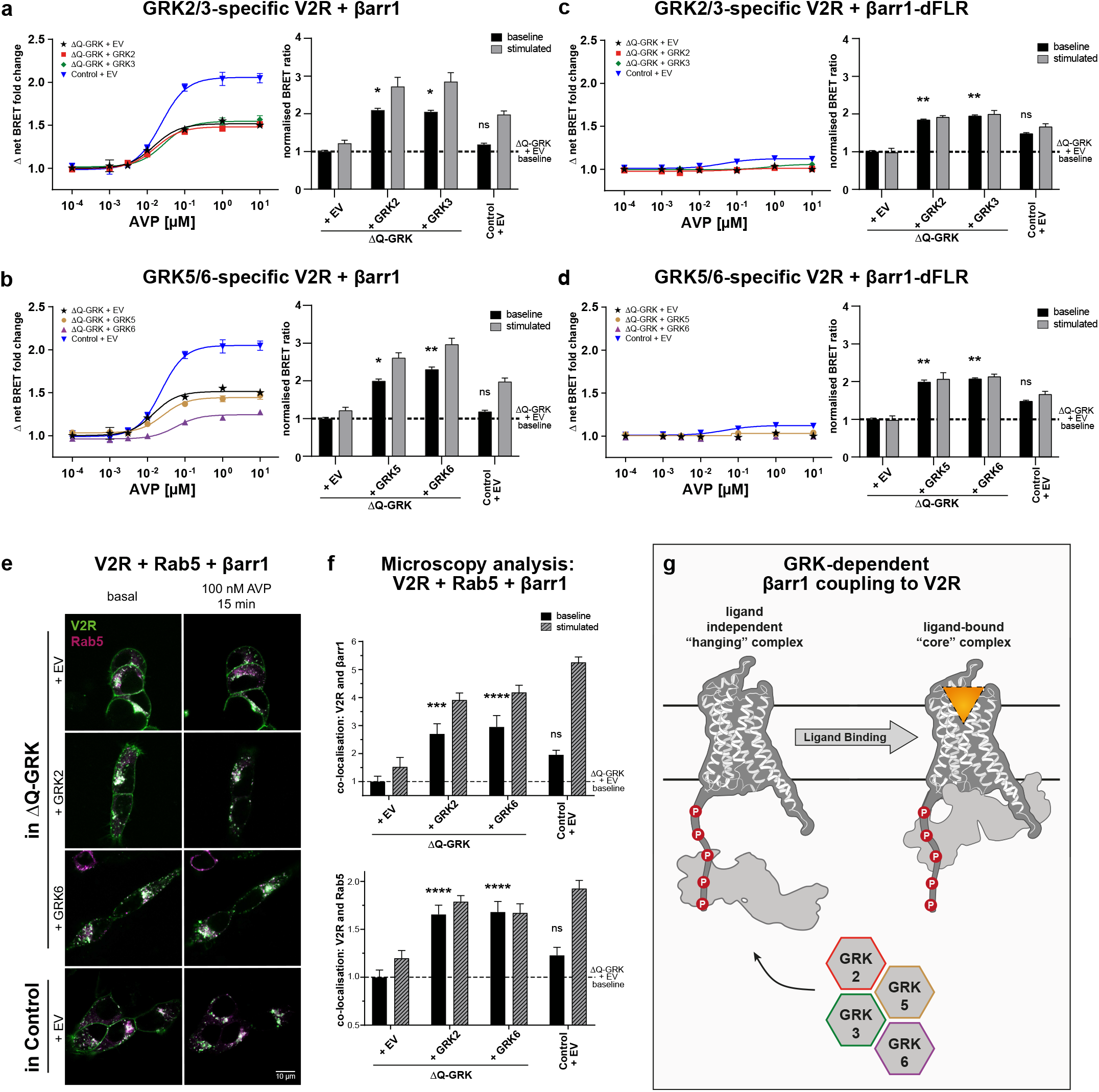
GRK2, 3, 5, or 6 can individually mediate a ligand-independent interaction of the V2R and β-arrestin1. **a-d** ΔQ-GRK or Control cells were transfected with V2R-Halo-Tag and one of the following β-arrestin1 (βarr1)-NanoLuc fusion constructs: wild type (**a, b**) or βarr1 lacking the finger loop region (dFLR; **c, d**). Additionally, either GRK2, 3, 5, 6 or the empty vector (EV) were transfected as indicated. The dynamic BRET changes are shown as ligand concentration-response curves normalised to baseline values and vehicle control. All data points are calculated as Δ net BRET fold change as the mean of three independent measurements ± SEM. The same dataset is presented as bar graphs, displaying the measured BRET-values before (baseline) and after stimulation with 10 μM [Arg^8^]-vasopressin (AVP; stimulated), normalised to the basal BRET ratio derived from the ΔQ-GRK + EV condition (dashed line). To test whether the baseline BRET ratios were significantly elevated compared to the respective ΔQ-GRK + EV baseline, an ANOVA and one-sided Dunnett’s test was performed (*p < 0.05; **p < 0.01; ns (not significant)). **e, f** ΔQ-GRK or Control cells were transfected with V2R-CFP (green), Rab5-mCherry (magenta), βarr1-YFP, and either EV, GRK2, or GRK6 as indicated. At least 30 images per condition, representing at least three independent transfections and experimental days, were taken before (basal) and after 15 minutes of 100 nM AVP stimulation. Representative images are shown in **(e)**, and **Supplementary Figure 6**. The co-localisation of V2R and βarr1 or Rab5 was quantified using Squassh and SquasshAnalyst **(f)**. Data are presented as mean fold change in co-localisation signal + SEM normalised to unstimulated (baseline) ΔQ-GRK + EV condition. Co-localisation prior to stimulation was compared using ANOVA and two-sided Dunnett’s test (* p < 0.05, ** p < 0.01, *** p < 0.001, ns (not significant)). **g** Schematic depiction of βarr1 interactions with the V2R in absence and presence of ligand, facilitated by high expression levels of GRKs.

The concentration-response curves shown to this point only reflect the fold change between the measured baseline and stimulated BRET ratios. The elevated baselines explain the unexpectedly low dynamic BRET changes, even though the absolute values of stimulated BRET ratios show a clear increase in the presence of overexpressed GRKs. Thus we conclude that already the basal molecular interaction between the V2R and β-arrestins is increased under those conditions.

Two major interaction interfaces between GPCRs and β-arrestins have been proposed. Namely the interaction mediated by the phosphorylated C-terminus of the receptor only (“hanging” complex)^15,31^, as well as the additional insertion of the arrestin finger loop region (FLR) into the intracellular cavity of the GPCR (“core” complex)^14^. Since we expect the unstimulated V2R to be in an inactive conformation, we hypothesise that the interaction between β-arrestins and the intracellular cavity of the GPCR is prevented. To test this hypothesis we measured β-arrestin recruitment with biosensors lacking the FLR (β-arrestin1/2-dFLR) **(Figure 3c, d; Supplementary Figure 6c, d)**.

For β-arrestin1-dFLR the baseline BRET measurements remained elevated, while agonist stimulation was not able to further increase the interaction between β-arrestin1-dFLR and the V2R **(Figure 3c, d)**. Thus we propose ligand-independent precoupling of β-arrestin1 to the V2R in a “hanging” complex in presence of overexpressed GRKs. The remaining ligand-dependent increase in β-arrestin1 recruitment might be explained by ligand-activation of the pre-coupled “hanging” complex and subsequent engagement of the FLR to form a tight “core” complex **(Figure 3g)**. In contrast, β-arrestin2 precoupling was found to depend on the FLR **(Supplementary Figure 6c, d)**.

Since the kinase-dependent precoupling of β-arrestins to the receptor was not observed for the chimeric β2V2 **(Supplementary Figure 4)**, we conclude that this effect must somehow be mediated by the transmembrane helix bundle.

Via confocal microscopy, we observed that the individual overexpression of GRK2 and 6 significantly increased the ligand-independent co-localisation between the V2R and Rab5 or β-arrestin1 in comparison to ΔQ-GRK+EV **(Figure 3e, f; Supplementary Figure 6e-h)**. This confirms that ligand-independent V2R–β-arrestin interactions, as facilitated by overexpressed GRKs, lead to functional receptor internalisation in line with Snyder *et al*.^32^.

### Distinct AT1R–β-arrestin complex configurations are mediated by GRK2/3, GRK5/6, or PKC

The assessment of β-arrestin complex configurations was found to be more intricate for the AT1R in comparison to the V2R. Hence, we arranged the data obtained for the AT1R in a kinase-specific manner in **Figure 4a-e (** GRK3 see **Supplementary Figure 7a)**. We observed the most prominent difference between angiotensin II (AngII)-induced β-arrestin1 and 2 recruitment to the AT1R, in ΔQ-GRK, as this condition features a pronounced higher recruitment of β-arrestin2 **(Supplementary Figure 3g)**. Since the AT1R is a known target of heterologous desensitisation^29^, we anticipated that PKC could be responsible for mediating this difference^33^. Hence, we conducted the experiment in the presence of Gö6983, a pan PKC inhibitor. This reduced the recruitment of both β-arrestins in Control and abolished GRK-independent β-arrestin1 recruitment **(Supplementary Figure 7b, c, 8a, b)**. In contrast, PKC inhibition had a negligible effect on the dynamic β-arrestin recruitment in the presence of overexpressed GRKs. Surprisingly, the overexpression of GRK5 and 6 showed lower dynamic recruitment of β-arrestin2 in comparison to ΔQ-GRK, regardless of PKC activity **(Figure 4de, e; Supplementary Figure 7b, c)**.

**Figure 4:**
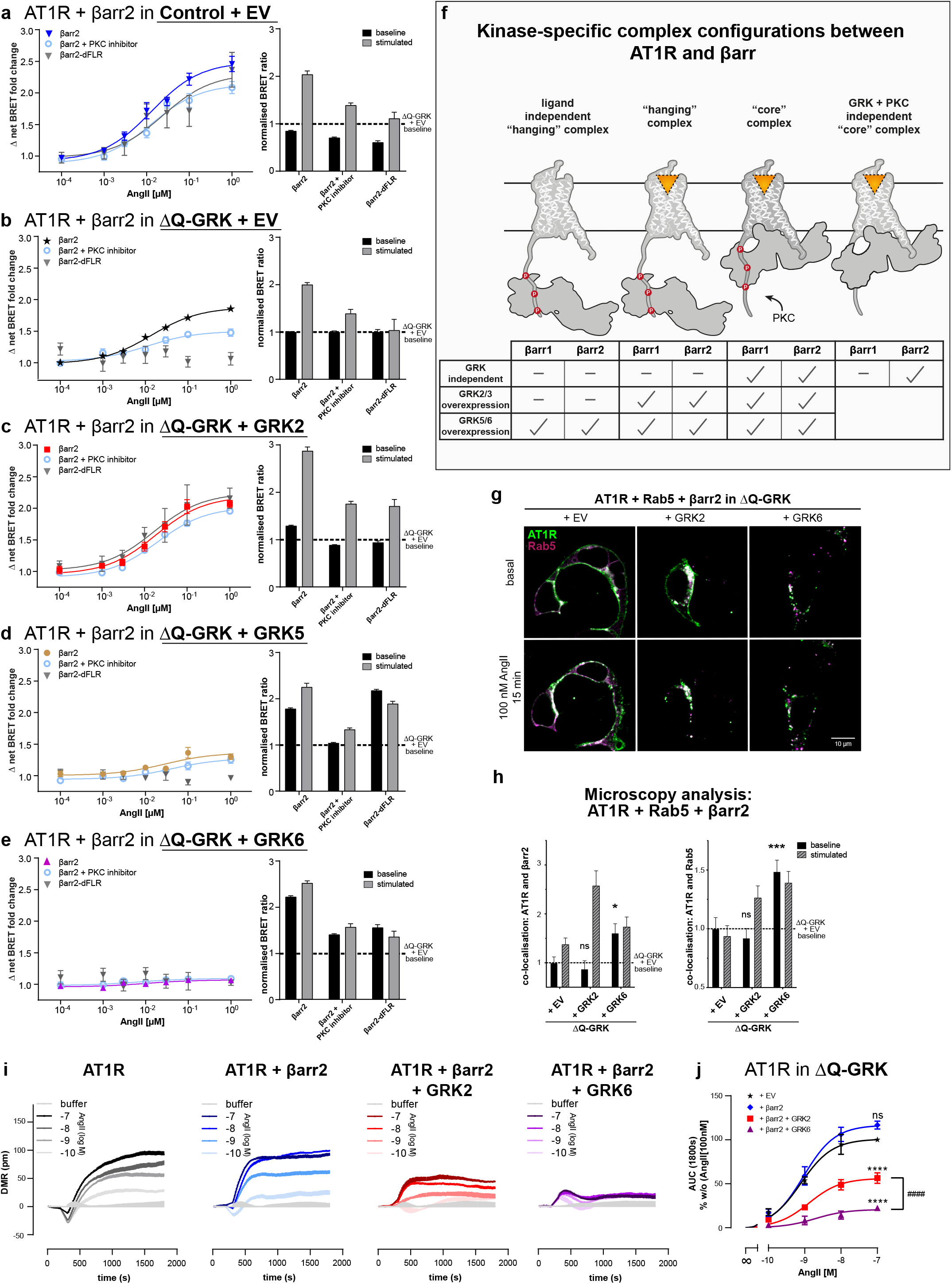
Distinct AT1R–β-arrestin2 complex configurations are mediated by GRK2/3, GRK5/6, or PKC. **a-e** To identify the contribution of the individual GRKs to the complex formation of β-arrestin2 (βarr2) with AT1R, Control or ΔQ-GRK cells were transfected with AT1R-NanoLuc, Halo-Tag-βarr2 and either GRK2, 3, 5, 6, or the empty vector (EV) as indicated, in absence or presence of PKC inhibitor Gö6983 (500 nM). Additionally, the GRK-specific βarr2 recruitment to the AT1R was measured utilising a Halo-Tag-βarr2 construct lacking the finger loop region (dFLR). Angiotensin II (AngII)-induced dynamic BRET changes are shown as concentration-response curves. All data points are calculated as Δ net BRET fold change normalised to baseline values and vehicle control, represented as the mean of at least three independent measurements ± SEM. The data are also presented in respective bar graphs, displaying the BRET ratios measured before (baseline) and after stimulation with 1 μM AngII (stimulated), normalised to the basal BRET ratio derived from the corresponding ΔQ-GRK + EV condition in **(b)**. **f** Schematic summary of the kinase-specific complex configurations between AT1R and βarr1 or 2 either in the absence of GRKs, mediated by GRK2/3 or GRK5/6 overexpression as observed in **(a-e; Supplementary Figure 7, 8)**. **g, h** ΔQ-GRK cells were transfected with hsp-AT1R-CFP (green), Rab5-mCherry (magenta), βarr2-YFP, and either EV, GRK2, or GRK6, as noted. At least 30 images per condition, representing at least three independent transfections and experimental days, were taken before (basal) and after 15 minutes of 100 nM AngII stimulation. Representative images are shown in **(g)**, and **Supplementary Figure 9**. The co-localisation of AT1R and βarr2 or Rab5 was quantified using Squassh and SquasshAnalyst **(h)**. Data are presented as mean fold change in co-localisation signal + SEM normalised to unstimulated ΔQ-GRK + EV. Co-localisation prior to stimulation was compared using ANOVA and two-sided Dunnett’s test (* p < 0.05, ** p < 0.01, *** p < 0.001, ns (not significant)). **i** ΔQ-GRK cells transiently transfected with hsp-AT1R-CFP, with or without co-transfection of either βarr2 alone or in combination with GRK2 or GRK6, were stimulated with AngII and real-time dynamic mass redistribution (DMR) responses were recorded as a measure of AT1R activity. DMR recordings are shown as mean + SEM of three technical replicates from a single experiment, representative of three to four independent experiments. **j** Concentration-effect curves derived from three to four independent experiments (representative shown in **i)** ± SEM are plotted from the area under the curve (AUC) within 0 and 1800 seconds. Data are normalised to the picometer wavelength shifts evoked with 100 nM AngII in cells expressing hsp-AT1R-CFP alone. Statistical significance was calculated using a two-way ANOVA (*p < 0.05; ** p < 0.01; *** p < 0.001; **** p < 0.0001; ns (not significant); #### p < 0.0001)

Again, analysis of the measured BRET ratios reveals significantly increased basal molecular interaction in the presence of overexpressed GRK5 and 6 **(Figure 4d, e; Supplementary Figure 4)**. This finding identifies the AT1R as another receptor that can interact with β-arrestins in a ligand-independent fashion, similar to the V2R. Interestingly, in this specific case GRK5 and 6 seem to be able to mediate these interactions. Notably, our statistical analysis shows a significant increase for the baseline of GRK3-mediated β-arrestin recruitment as well. However, since the greater part of the recruitment seems to be ligand-dependent in this condition, we did not conclude efficient AT1R–β-arrestin precoupling mediated by GRK3.

After disclosure of these fundamental kinase-specific effects, we investigated whether these AT1R–β-arrestin complexes occur in a “hanging” or “core” configuration **(Figure 4; Supplementary Table 3)**. As stated above, PKC inhibition reduced β-arrestin2 recruitment in ΔQ-GRK, but the recruitment is still detectable. The deletion of the FLR abolished β-arrestin2 recruitment independently of PKC activity **(Figure 4b)**. Thus we conclude that the FLR is essential for the PKC-mediated AT1R–β-arrestin2 complex, as well as a GRK- and PKC-independent complex. This suggests that PKC activity alone cannot mediate a “hanging” complex configuration.

In the presence of overexpressed GRK2 or 3, dynamic β-arrestin2 recruitment is neither altered by PKC inhibition nor deletion of the FLR **(Figure 4c; Supplementary Figure 7a)**. Interestingly, upon PKC inhibition the measured absolute BRET ratios are decreased to the levels recorded for the dFLR construct. This led us to the assumption that PKC inhibition and the deletion of the FLR mediate the same biological effect. Since PKC activity alone was shown to only mediate a “core” complex configuration **(Figure 4b)** and the FLR is dispensable for GRK2/3-mediated β-arrestin2 binding, we propose that GRK2 and 3 predominantly facilitate the formation of a “hanging” complex. Despite this, with our experimental setup, we cannot exclude the formation of a “core” complex between the two proteins under physiological conditions with unaltered PKC activity.

The precoupling effect mediated by GRK5 and 6 **(Figure 4d, e)** is also observed for the β-arrestin2-dFLR mutant, suggesting the formation of a ligand-independent “hanging” complex with β-arrestin2. Again, the remaining ligand-dependent increase in β-arrestin2 recruitment measured for GRK5 overexpression could reflect on ligand-activation of the pre-coupled “hanging” complex and subsequent formation of a tight “core” complex **(Figure 4f)**, similar to the mechanism proposed for the V2R **(Figure 3g)**. Notably, GRK6-mediated precoupling was reduced, indicating that the FLR plays a role in this process.

The utilisation of kinase dead (KD) mutants revealed that the precoupling of β-arrestin2 is mediated by GRK5 or 6 kinase activity **(Supplementary Figure 7d, f, h)**. Particularly, the phosphorylation of inactive AT1R in the presence of overexpressed GRK5 has been reported before^34^. Interestingly this β-arrestin2 precoupling effect was not observed in ΔGRK2/3/6 or ΔGRK2/3/5 **(Supplementary Figure 7e, g)**, indicating that it is dependent on individual GRK expression levels. This could explain how the same receptor might be differentially regulated in specific tissues, cellular compartments^8^ or under pathophysiological conditions featuring dysregulated GRK expression levels^35^.

In case of β-arrestin1, GRK5 and 6 overexpression also led to an enhanced basal interaction with AT1R **(Supplementary Figure 8g, h)**. In general, the GRK-dependent interaction of both β-arrestin isoforms and the AT1R are similar. Additionally, we were able to conclude that both β-arrestins can use PKC phosphorylation to stabilise a “core” complex with the AT1R. Interestingly, we observed recruitment of β-arrestin2 in absence of GRK and PKC phosphorylation, whereas β-arrestin1 does not seem to be able to interact with the unphosphorylated receptor. Thus we can exclude the formation of a GRK- and PKC-independent “core” complex for β-arrestin1 **(Figure 4f, Supplementary Figure 8)**.

### Ligand-independent AT1R regulation by GRK6 leads to receptor internalisation and impaired signalling responses

To test if these different kinase effects have a direct impact on receptor functionality, we employed confocal microscopy and dynamic mass redistribution (DMR) measurements. In ΔQ-GRK the AT1R did not show pronounced internalisation, while GRK2 overexpression strongly supports AngII-dependent receptor internalisation. In contrast, the AT1R was already found in intracellular compartments when overexpressing GRK6, independently of ligand application **(Figure 4g, h; Supplementary Figure 9)**.

As expected, AngII evoked robust primary receptor signalling in GRK-deficient cells when assessed by DMR **(Figure 4i, j)**. Interestingly, co-transfection of ΔQ-GRK with β-arrestin2 did not suffice to diminish cellular AT1R signalling despite significant β-arrestin2 recruitment under comparable experimental conditions **(Figure 4i** compare with **4b)**. Even though PKC-specific phosphorylation can stabilise AT1R–β-arrestin2 interactions **(Figure 4b)**, we conclude that GRK phosphorylation is strictly required for β-arrestin2-mediated receptor desensitisation and internalisation **(Figure 4j)**. The cellular signalling responses were significantly dampened under GRK2 overexpression, confirming efficient receptor desensitisation in this condition. In this system GRK2 functions as a canonical sensor for receptor activation, governing location and activity of GPCRs via mediation of β-arrestin functions.

The overexpression of GRK6 almost eliminated the measured cellular signalling response **(Figure 4i)**. This, in combination with the observed pattern of AT1R subcellular localisation, suggests that GRK6-mediated AT1R phosphorylation precludes a majority of receptor molecules from the membrane and thus from being ligand-activated. This is especially significant, as it demonstrates that the upregulation of GRKs could have two distinctly different consequences. Depending on the combination of involved kinases and receptors, it could either lead to a canonical increase in efficiency to induce arrestin-mediated receptor regulation or an almost complete loss of GPCR responsiveness. Notably, we cannot exclude that internalised receptors can still signal from intracellular compartments.

## Discussion

Using our triple GRK KO cell lines, it became evident that different GPCRs require certain levels of GRK expression in order to recruit arrestins. In contrast to the β2ADR **(Figure 1c)**, the PTH1R and M5R showed robust β-arrestin2 recruitment in ΔGRK3/5/6 cells **(Supplementary Figure 5)**. This does not reflect on the ability of GRK2 to regulate the β2ADR, as has been shown multiple times in literature and our presented overexpression experiments **(Figure 1e)**. These results demonstrate that the affinities of GRK isoforms to GPCRs differ depending on the individual receptor. Thus, the tissue-specific expression levels of individual GRKs in combination with their affinities to specific receptors determine GPCR regulation.

Several studies have demonstrated phosphorylation of the β2ADR at different serine and/or threonine residues^36,37^ and phosphorylation by GRK2 or 6 was shown to serve different functions^7^. In our study we observed that all GRKs can mediate receptor–β-arrestin interactions to the same extent and by using a single β-arrestin2 conformational change sensor (FlAsH5 according to Nuber *et al.* 2016^27^), we could demonstrate that the N-domain of β-arrestin2, which recognises the phosphorylated receptor C-terminus, showed similar conformational changes for the different kinases **(Figure 1g)**. However, more experiments have to be performed to rule out that differential β2ADR phosphorylation by GRK2 or 6 might lead to distinct β-arrestin2 conformational changes. While different GRK isoforms might preferably phosphorylate distinct sites of the β2ADR, resulting in different C-terminal phosphorylation patterns, our experiments clarify that the phosphorylation by each GRK isoform is sufficient to induce high-affinity β-arrestin recruitment **(Figure 1e)**.

Interactions between arrestins and the M2R have investigated previously^38–40^. Interestingly, the β-arrestin recruitment assay only induced minimal BRET changes for the M2R at endogenous GRK expression levels **(Supplementary Figure 10)**. However, upon overexpression of GRK2 or 3 robust β-arrestin recruitment was observed. It is tempting to speculate that the M2R exhibits a rather low affinity for GRKs to prevent its desentitisation since its function is essential for the reduction of heart rate^41^. Under pathophysiological conditions of GRK2 overexpression during chronic heart failure^42^, the M2R might internalise which could possibly contribute to tachycardiac effects in patients.

Strikingly, we were able to show that certain GPCRs are readily being regulated by overexpressed GRKs in a cellular system without ligand addition. Although, ligand-independent GPCR phosphorylation has been described for multiple receptors^34,37,43–45^, it has not been convincingly shown to this point that this phosphorylation would translate into arrestin functions in a cellular context. As seen for the MOP, GRK5/6-dependent phosphorylation does not result in arrestin-dependent receptor regulation **(Supplementary Figure 1d-g)**. We demonstrated that both, the V2R and AT1R couple to arrestins and internalise in a ligand-independent fashion as long as the essential GRKs are present at high expression levels **(Figure 3 and 4)**. Although GRK5 and 6 mediate this β-arrestin precoupling for both receptors, this is not a unique feature of the membrane-associated GRK4 family kinases. As GRK2 and 3 can achieve similar effects for the V2R, this rather has to be a distinct characteristic of a specific GPCR. Especially, since this precoupling effect could not be transferred to the β2ADR by exchange with the V2R C-terminus **(Supplementary Figure 3b, c, m, 4)**, we can also exclude the C-terminus as sole mediator of GRK-specific processes.

Currently it is unknown whether a single receptor is phosphorylated by a single kinase or by multiple kinases in a sequential manner. This could lead to vastly different outcomes, as we were able to show that different GRKs and second messenger kinases are able to induce divergent regulatory processes, depending on the targeted GPCR. Notably, the effect of PKC on the regulation of GRKs was described multiple times in literature^46–48^, as it was shown that the activity of GRKs can be either increased or decreased via PKC activity, depending on the used system. Even though, we were able to record a reduction in GRK5- and 6-mediated β-arrestin recruitment to the AT1R under inhibition of PKC **(Figure 4d, e)**, we could not draw a conclusion on how PKC modulates the activity specific kinases. The elucidation of these intriguing effects require more in depth analysis and possibly single molecule studies.

For each receptor case with unclear GRK assignment by statistical analysis **(Figure 2i, j)**, alternative kinases were previously reported to be involved in receptor phosphorylation. In case of the M1R and M3R, casein kinase 1 alpha and casein kinase 2 were shown to be involved in receptor phosphorylation, respectively^49,50^. For the C5aR1, PKCβ was shown to contribute to receptor phosphorylation^51^. Therefore, our cellular platform for arrestin recruitment might be able to rapidly differentiate between receptors with purely GRK-dependent arrestin recruitment and receptors that rely on the action of other intracellular kinases for efficient arrestin-binding, desensitisation, and internalisation.

Using our ΔQ-GRK cell line, we were able to show that high expression levels of GRK2 and 3 are able to mediate β-arrestin interactions with all tested GPCRs. GRK5 and 6 seem to fulfil divergent roles depending on the analysed GPCR, as they were not able to induce β-arrestin-coupling to the M2R, M4R, M5R and MOP **(Figure 2i, j; Supplementary Figure 3)**. Interestingly, this is not necessarily due to a lack of receptor phosphorylation, as we were able to show that GRK5 and 6 phosphorylate the MOP upon agonist activation, but fail to mediate β-arrestin recruitment and receptor internalisation **(Supplementary Figure 1)**. Nevertheless, we found receptors for which the GRK5- and 6-facilitated receptor regulation is indistinguishable from that mediated by GRK2 and 3 (namely the PTH1R and β2ADR, **Figure 1 and 2**).

In an endeavour to match the measured GRK-specific β-arrestin recruitment with the main features of the tested GPCRs, we analysed the length of the respective C-terminus and intracellular loop 3 (IL3), as well as the number and relative location of their putative phosphorylation sites **(Supplementary Table 1, Supplementary Figure 11)**. Here we were not able to find correlations that would compellingly explain the observed GRK-selectivity for our panel of GPCRs **(Supplementary Figure 11a)**. Interestingly, none of the analysed class B receptors are selectively regulated by GRK2 and 3 only ( β2V2 was excluded from this analysis as an unphysiological chimera; **Supplementary Figure 11b)**.

Further, we assessed the abundance and relative positions of previously established phosphorylation motifs (PPP, PXPP, PXPXXP, PXXPXXP, whereas P represents either Ser, Thr, Asp, or Glu residues and X any amino acid), reported to be important for β-arrestin recruitment^52,53^. As illustrated in **Supplementary Figure 11c-n**, all motifs show a similar distribution between our defined GRK-selectivity groups.

Interestingly, the PXPP motif showed a higher abundance in the central area (0.25 – 0.75) of analysed peptide stretches of GPCRs regulated by GRK2/3. In contrast, the same motif was found more often in the peripheral area (0.00 – 0.25 and 0.75 – 1.00) of peptide stretches of GPCRs regulated by GRK2/3/5/6 **(Supplementary Figure 11h)**. This is the only instance with a significant association (Fisher’s exact test, p=0.0005367) between position of putative phosphorylation motifs (PXPP: central vs peripheral) and GRK-specificity of the assessed GPCRs, according to our analysis. Notably, although class A and B GPCRs are differentially represented in the GRK-specificity groups as defined in **Figure 2i, j**, no significant association was found between the positions of PXPP motifs and the A-B classification **(Supplementary Figure 11l;** Fisher’s exact test, p=0.7328). However, the causality of this association remains to be explained. In general, we did not identify common features of C-terminal and IL3 sequences which would allow for the reliable prediction of GRK-selectivity. Hence, we propose that GRK-selectivity is rather defined by the overall geometry of each GPCR and influenced not only by the availability of putative phosphorylation sites, but also by the general promiscuity of other intracellular domains.

While GRK5 and 6 are membrane-localised^54^, GRK2 and 3 are primarily cytosolic and translocate to the plasma membrane supported by interactions with βγ-subunits of activated G proteins^55^. It is still conceivable that GRK5 and 6 interactions with certain GPCRs might be obstructed due to their distinct cellular localisation. The plasma membrane features a rather heterogeneous distribution of proteins and it has been shown that certain GPCRs tend to reside in specific membrane confinements^56^. Some receptors might localise in membranous compartments that are inaccessible for GRK5 and 6. Following this hypothesis, these GPCRs would be accessible to GRK2 and 3 since they emerge from the cytosol and would not be limited to two dimensional diffusion and hindered by possible confinements. This still does not exclude the existence of GPCRs which do not serve as substrates for GRK2 or 3, due to e.g. low affinity. In conclusion, we were able to elucidate the GRK-specificity of receptor regulation for 12 different GPCRs. Our analysis demonstrates that different GRK isoforms may have identical, overlapping or divergent functions, depending on the targeted GPCR. This adds another layer of complexity to the regulation of GPCR signalling and trafficking and a possible explanation of how different β-arrestin functions are mediated across various tissues and cell types, especially considering often dysregulated, pathophysiological GRK expression levels.

## Acknowledgements

We want to thank Ulrike Schiemenz and Nina Kathleen Blum for assistance with the MOP-internalisation studies, Dr. Aurélien Rizk for the help in microscopy data analysis, Prof. Tom Kirchhausen for providing the Rab5-mCherry plasmid, and the Core Facility Flow cytometry of the FLI – Leibniz Institute for Age Research, Jena, for sorting of the stable cell lines.

## Author contributions

J.D. engineered all GRK knockout cells; J.D., R.S.H, C.H. developed the concept and designed the experiments; J.D., R.S.H., E.S.F.M., M.R., S.B, C.Z., L.K. conducted the experimental work; J.D., R.S.H., E.S.F.M., M.R. compiled the data; J.Z. designed, conducted, and analysed AT1R targeted DMR measurements, supervised by E.K.; S.F., A.K. planned, conduced, and analysed the MOP internalisation experiments with support from E.M.; S.S., C.H. supervised the project; S.S. provided phosphosite specific antibodies; J.D., R.S.H., E.S.F.M., M.R., C.H. wrote the manuscript; all other authors critically revised the manuscript and gave final approval.

---

**Supplementary Figure 1:**
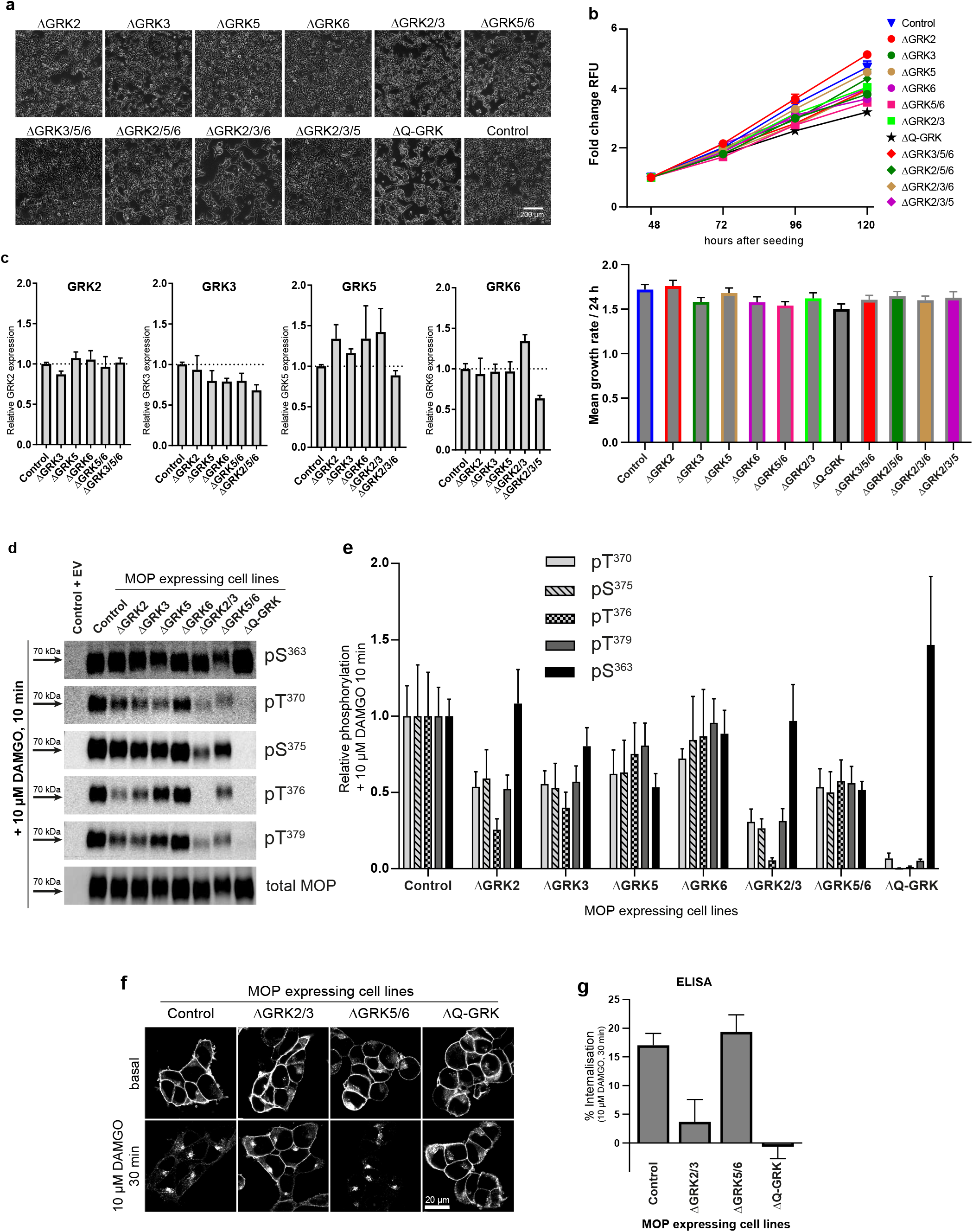
**a** Single (ΔGRK2, 3, 5, 6), double (ΔGRK2/3 or 5/6), triple (ΔGRK3/5/6, 2/5/6, 2/3/6, 2/3/5), and quadruple (ΔQ-GRK) GRK knockout cells were generated using the CRISPR/Cas9 technology and established as single cell clones. Cell morphology was documented by phase contrast microscopy. **b** Cell viability was determined by adding CellTiter-Blue reagent (Promega) at indicated timepoints and measuring relative fluorescence units (RFU) after 90 minutes of incubation. Shown are the respective growth curves as mean ± SEM of three independent experiments, expressed as fold change compared to 48 h after seeding (top panel). Growth rates per 24 h were calculated for each cell clone and are presented in bar graphs as mean + SEM (bottom panel). **c**. Quantification of the untargeted GRKs was performed by analysing western blots (representative blot in **Figure 1a**) made from four independent protein lysates of each cell clone and is depicted in bar graphs as mean + SEM, normalised to the respective remaining expression in Control cells. **d-g** ΔGRK clones and Control cells stably expressing an N-terminally HA-tagged μ-opioid receptor (MOP) were used for analysis of site-specific phosphorylation and receptor internalisation. **d** Prior to immunoprecipitation, cells were treated with the MOP agonist [D-Ala^2^, N-MePhe^4^; Gly-ol]-enkephalin (DAMGO) as indicated and subsequently analysed for site-specific phosphorylation by western blot. Shown are representative blots of four independent experiments. **e** Blots were quantified and relative phosphorylation compared to the respective Control signal of the indicated GRK target sites is shown as mean + SEM. **f** ΔGRK MOP cells were stimulated with DAMGO as indicated or left untreated (basal). After fixation, the cells were stained with anti-HA antibody followed by Alexa488-conjugated secondary antibody, and examined by confocal microscopy. Shown are representative images of five independent experiments. **g** Receptor internalisation was measured by ELISA. Data represent per cent loss of cell-surface receptors in DAMGO treated cells as compared to vehicle stimulated sister cultures. Data are presented as mean + SEM from at least six independent experiments.

**Supplementary Figure 2:**
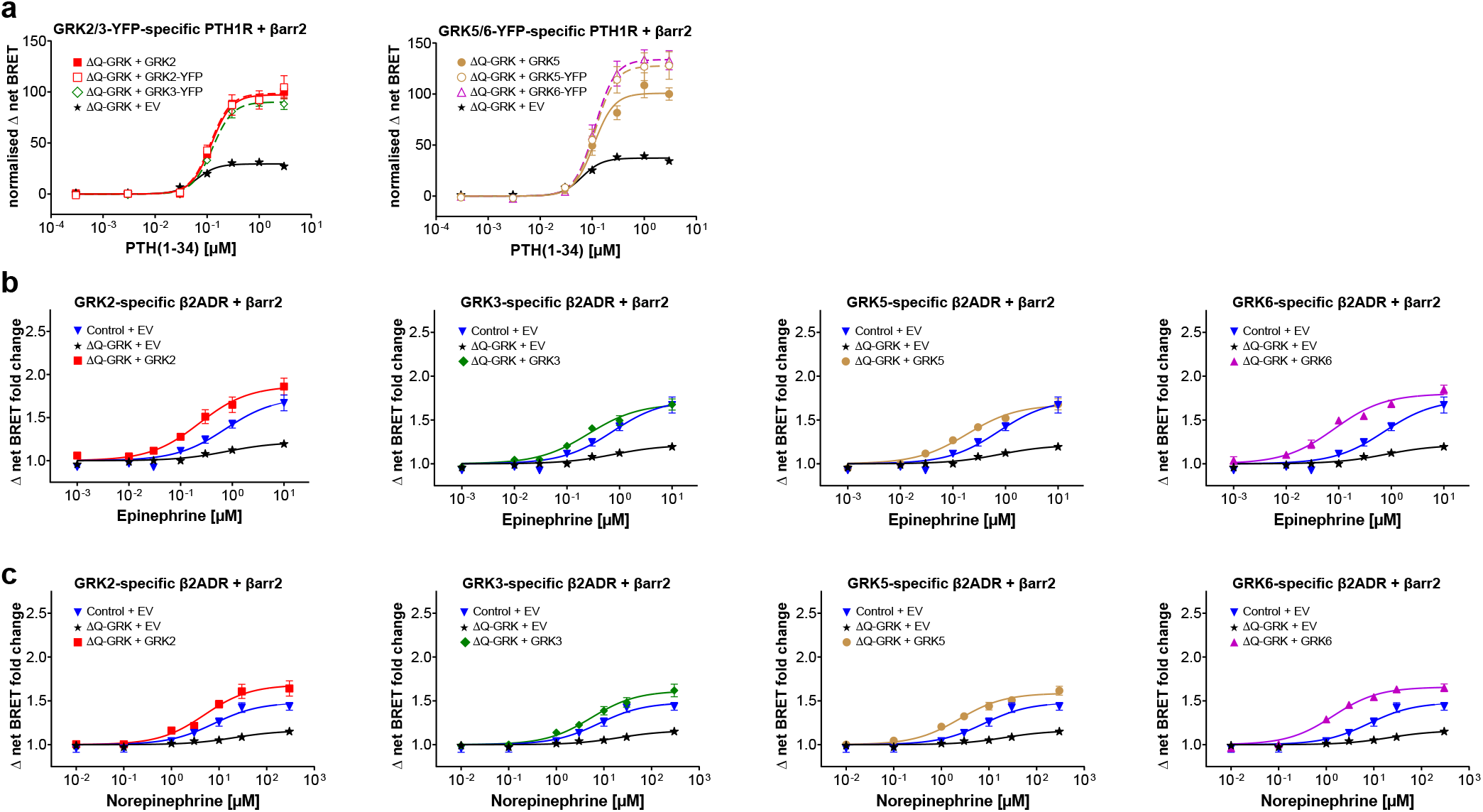
**a** ΔQ-GRK cells were transfected with either YFP-tagged or untagged GRKs as indicated and β-arrestin2 (βarr2) recruitment to the parathyroid hormone 1 receptor (PTH1R) was measured using our NanoBRET βarr recruitment assay. Data points are presented as the mean ± SEM of the calculated Δ net BRET fold change from three independent measurements. The utilisation of the GRK-YFP constructs resulted in comparable βarr2 recruitment as in presence of the untagged GRKs (corresponds to **Figure 1f**). **b, c** GRK-specific βarr2 recruitment to the β2ADR was measured in presence of the endogenous ligands epinephrine **(b)** or norepinephrine **(c)** (correspond to **Figure 1 i and j,** respectively). Individual GRK isoforms were overexpressed in ΔQ-GRK as indicated. For better comparability, the data for the Control + empty vector (EV) and ΔQ-GRK + EV conditions are depicted in each panel. The mean ± SEM of the Δ net BRET fold change from three independent experiments are shown.

**Supplementary Figure 3:**
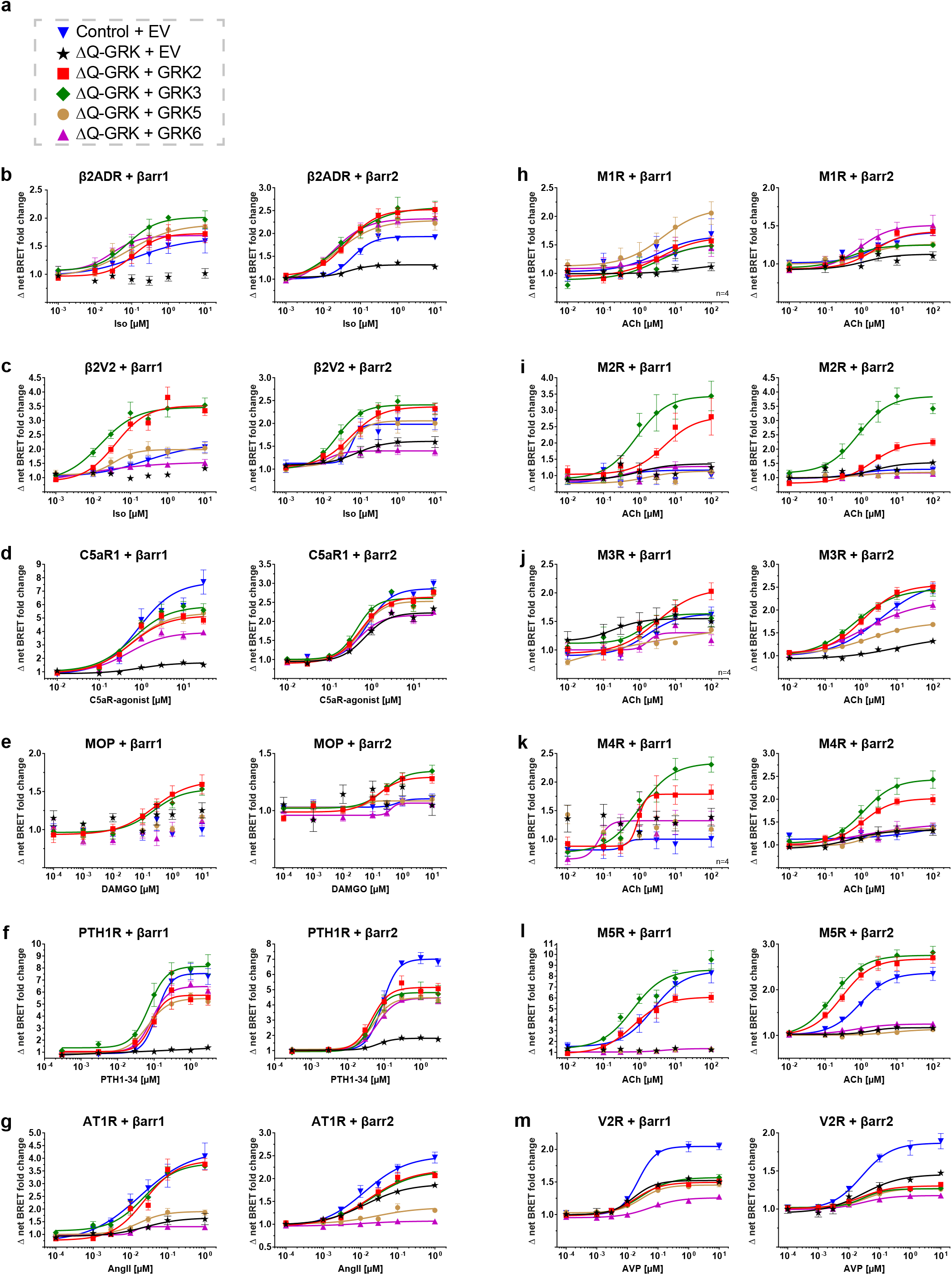
**a** Legend for concentration-response curves shown in **(b-m)**. **b-m** The Halo-Tag-β-arrestin (βarr) fusion protein is recruited to a Nano luciferase (NanoLuc)-tagged GPCR upon agonist activation and subsequent receptor phosphorylation. For the PTH1R and the V2R, the BRET pair was swapped. Overexpression of single GRKs in ΔQ-GRK cells allows the assessment of the impact of individual GRKs on this process. ΔQ-GRK or Control cells were transfected with the respective tagged GPCR and βarr1 or 2-fusion constructs. Additionally, either GRK2, 3, 5, 6, or empty vector (EV) were co-transfected as indicated. The dynamic BRET changes are shown as ligand concentration-response curves normalised to baseline values and vehicle control. All data points are calculated as Δ net BRET fold change as the mean of at least three independent measurements ± SEM. Results of the statistical analysis are listed in **Supplementary Table 2** and plotted in **Figure 2i**.

**Supplementary Figure 4:**
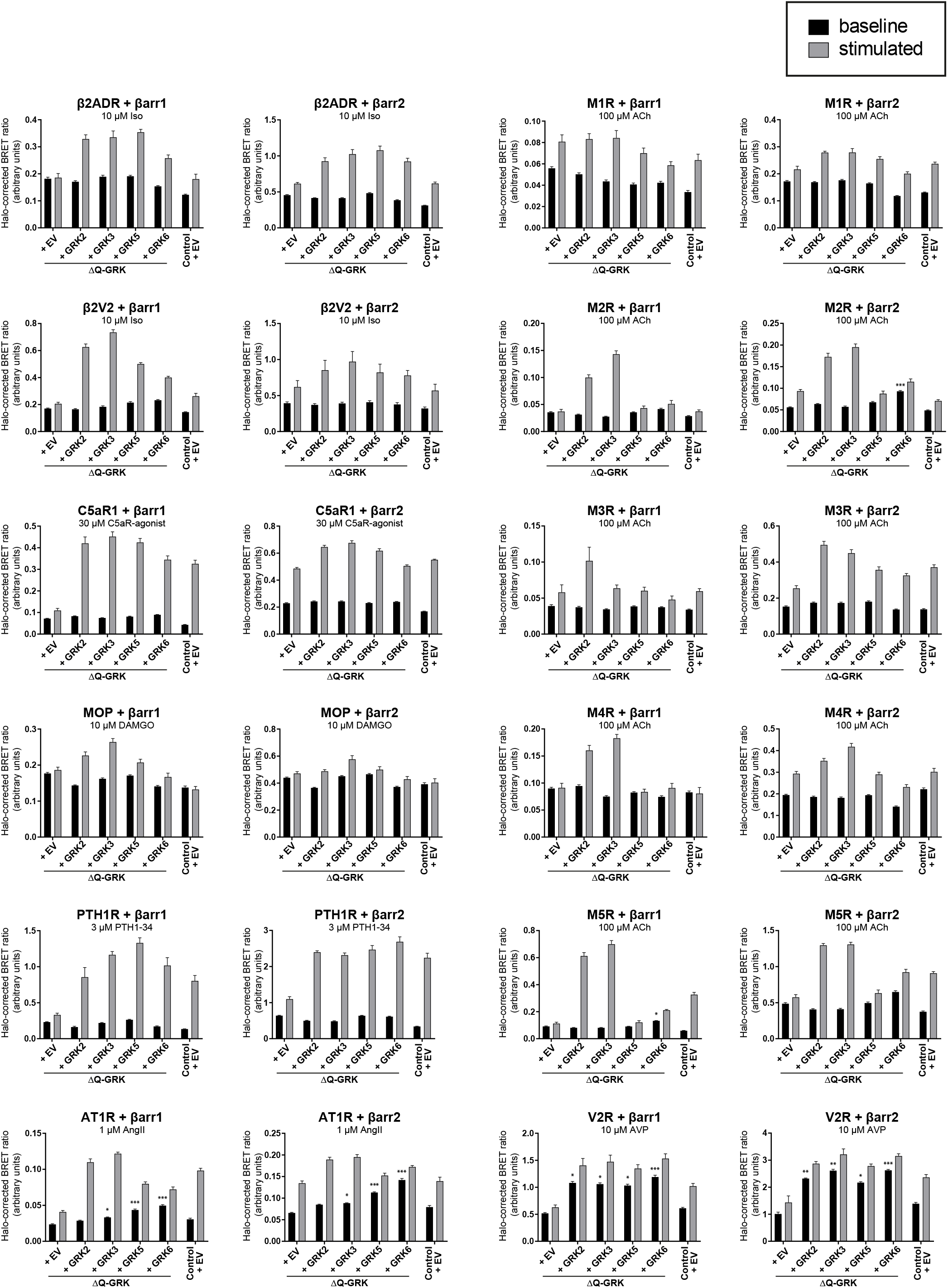
The data shown are derived from concentration-response curves in **Supplementary Figure 3**. ΔQ-GRK or Control cells were transfected with the Nano luciferase (NanoLuc)-tagged GPCR and Halo-Tag-β-arrestin (βarr)1 or 2 fusion constructs. For the PTH1R and the V2R, the BRET pair was swapped. Additionally, either GRK2, 3, 5, 6, or the empty vector (EV) were co-transfected. The non-normalised, Halo-corrected BRET ratio is presented before (baseline) and after stimulation with saturating agonist concentrations (stimulated) as indicated. The bar graphs show the mean of at least three independent measurements + SEM. To test whether the baseline BRET ratios were significantly elevated compared to the respective ΔQ-GRK + EV baseline, an ANOVA and one-sided Dunnett’s test was performed (* p < 0.05; ** p < 0.01; *** p < 0.001). Baselines without indication of significance were found to be not significant.

**Supplementary Figure 5:**
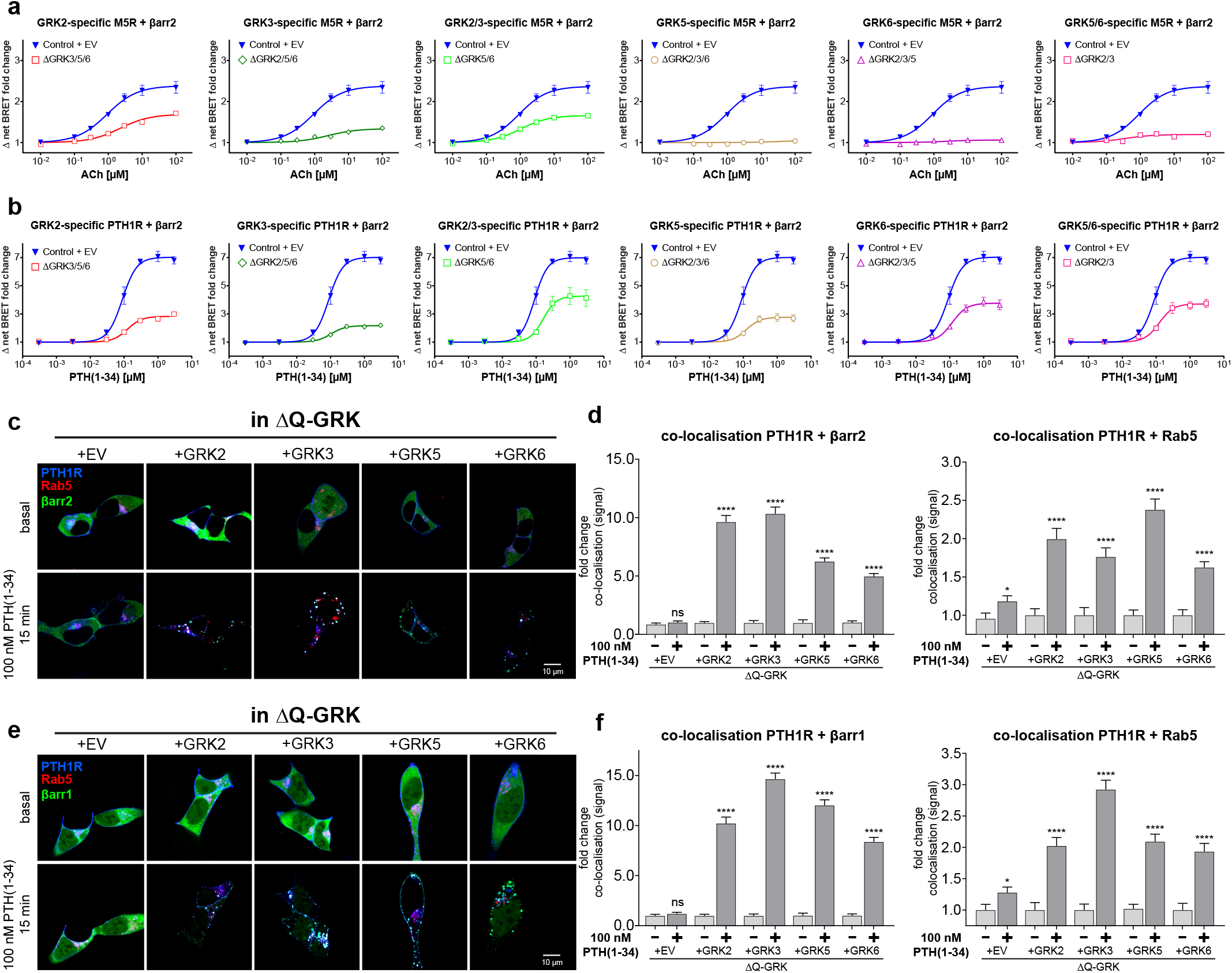
**a, b** β-arrestin2 (βarr2) recruitment to the muscarinic M5 acetylcholine receptor (M5R) **(a)** or parathyroid hormone 1 receptor (PTH1R) **(b)** in cells expressing all endogenous GRKs (Control + EV), only one remaining endogenous GRK (ΔGRK3/5/6, 2/5/6, 2/3/6, 2/3/5), or two remaining endogenous GRKs (ΔGRK5/6, 2/3) as indicated is shown as Δ net BRET fold change (mean ± SEM of three independent experiments). For easier comparison, Control curves are depicted in every panel. **c-f** Confocal live-cell microscopy was performed using ΔQ-GRK cells transfected with PTH1R-CFP (blue), Rab5-mCherry (red), βarr2-YFP **(c)** or βarr1-YFP **(e)** expression constructs (green) and individual untagged GRK2, 3, 5, or 6. Shown are representative images, taken before (basal) and after 15 minutes of stimulation with 100 nM parathyroid hormone (1-34) (PTH(1-34)). The normalised co-localisation of PTH1R and βarr2 or Rab5 **(d)** and βarr1 or Rab5 **(f)** was quantified using Squassh and SquasshAnalyst, with at least 30 images per condition. Data are presented as mean fold change in co-localisation signal + SEM. Statistical analysis was performed using a two-way mixed-model ANOVA followed by a paired t-test (* p < 0.05; ** p < 0.01; *** p < 0.001; **** p < 0.0001; ns (not significant)).

**Supplementary Figure 6:**
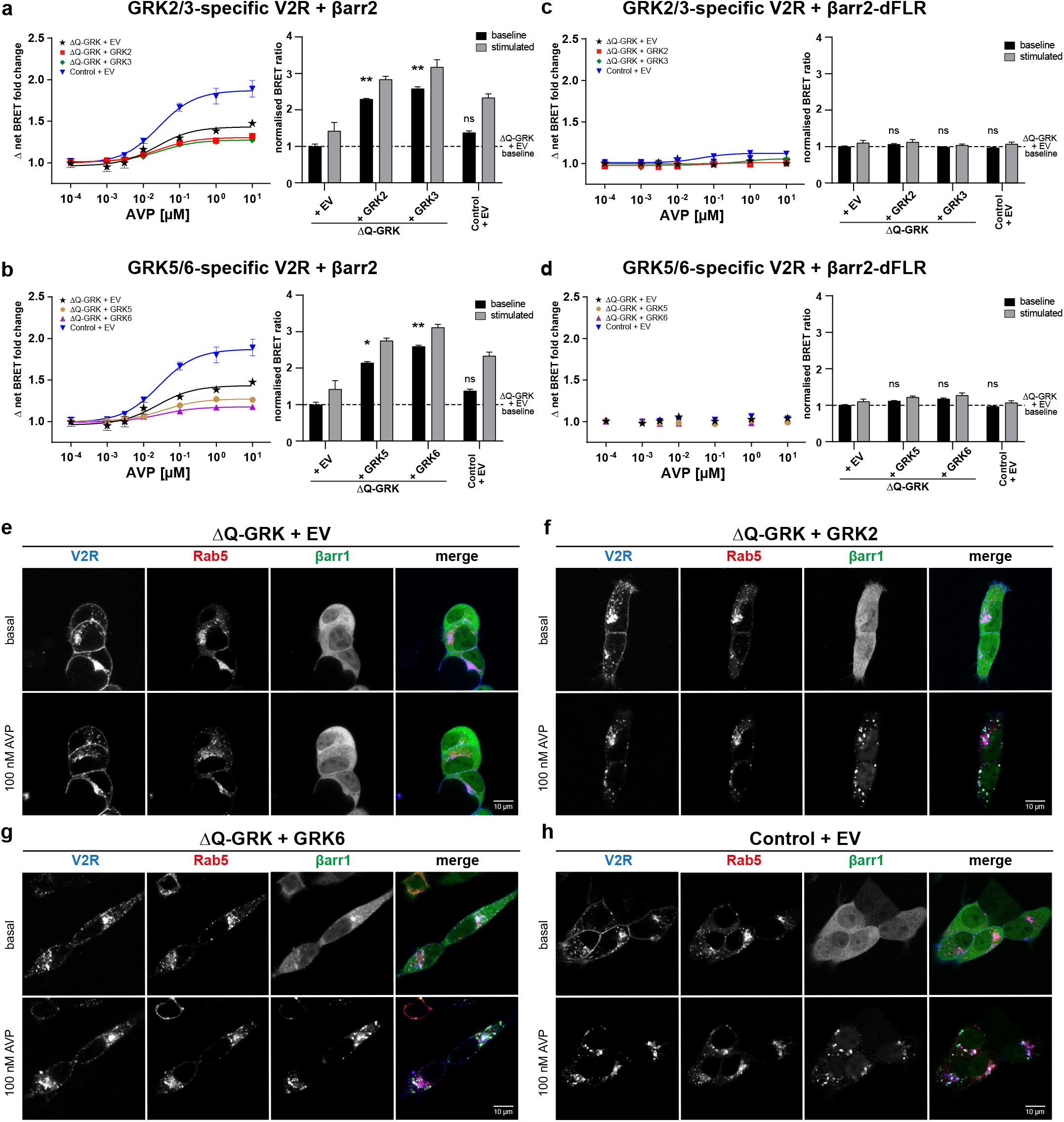
**a-d** ΔQ-GRK or Control cells were transfected with vasopressin 2 receptor (V2R)-Halo-Tag and one of the following β-arrestin(βarr) 2-Nano luciferase (NanoLuc) fusion constructs: wild type **(a, b)** or βarr2 lacking the finger loop region (dFLR; **c, d**). Additionally, either GRK2, 3, 5, 6 or the empty vector (EV) were transfected as indicated. The dynamic BRET changes are shown as ligand concentration-response curves normalised to baseline values and vehicle control. All data points are calculated as Δ net BRET fold change as the mean of three independent measurements ± SEM. The same dataset is presented as bar graphs, displaying the measured BRET-values before (baseline) and after stimulation with 10 μM [Arg^8^]-vasopressin (AVP; stimulated), normalised to the basal BRET ratio derived from the ΔQ-GRK + EV condition (dashed line). To test whether the baseline BRET ratios were significantly elevated compared to the respective ΔQ-GRK + EV baseline, an ANOVA and one-sided Dunnett’s test was performed (*p < 0.05; **p < 0.01; ns (not significant)). **e-h** (Corresponds to **Figure 3e**) ΔQ-GRK or Control cells were transfected with V2R-CFP (blue), Rab5-mCherry (red), βarr1-YFP (green), and either EV, GRK2, or GRK6 as indicated. At least 30 images per condition were taken before (basal) and after 15 minutes of 100 nM AVP stimulation. Single channels and overlay of all three channels from the corresponding images shown in **Figure 3 e** are depicted here.

**Supplementary Figure 7:**
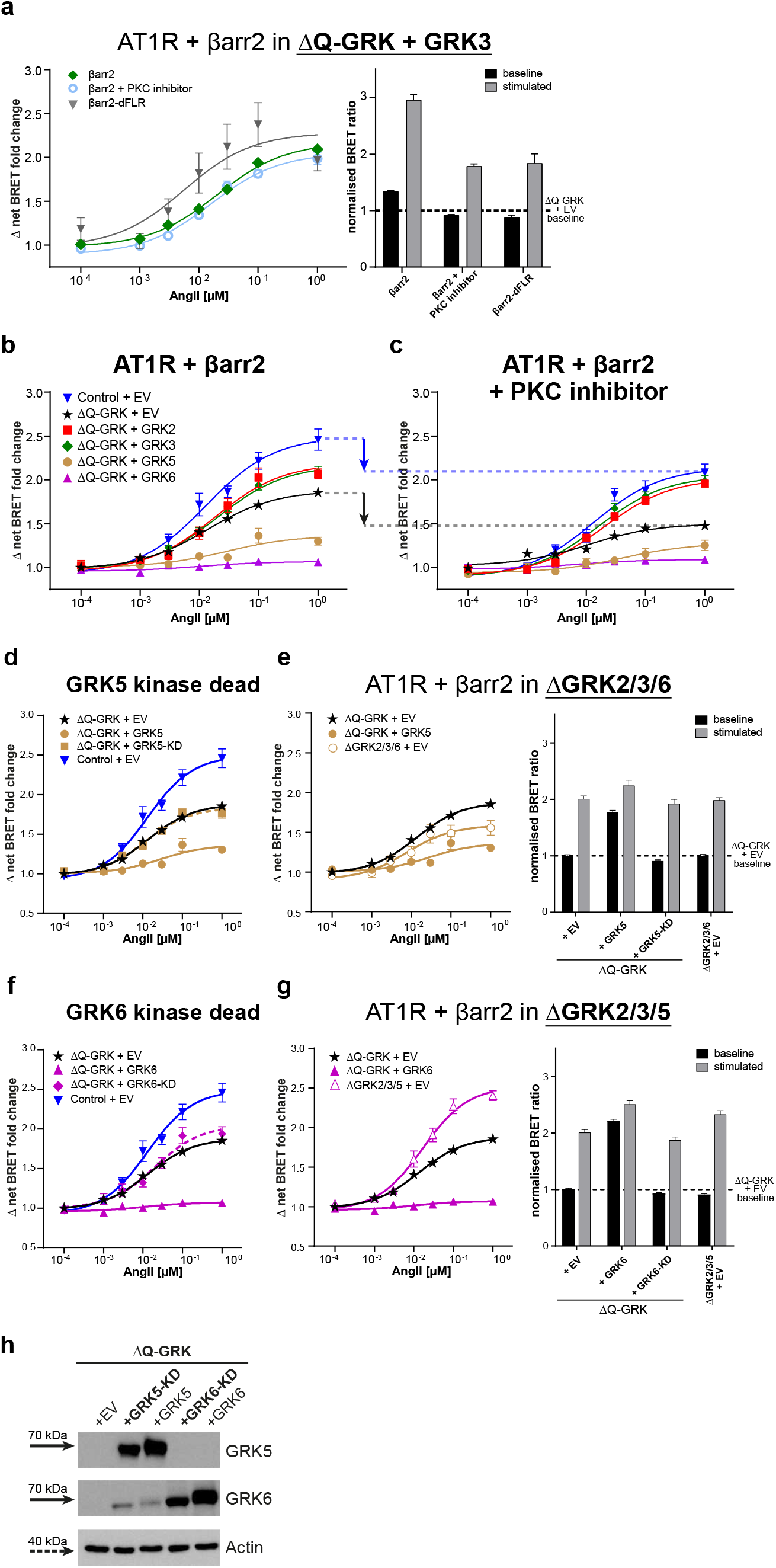
**a** Data for GRK3 are shown analogous to **Figure 4 a-e**. **b, c** ΔQ-GRK cells were transfected with Angiotensin 1 receptor (AT1R)-Nano luciferase (NanoLuc), Halo-Tag-β-arrestin2 (βarr2), and either GRKs or the empty vector (EV) as indicated. βarr2 recruitment to the AT1R was measured in absence or presence of PKC inhibitor Gö6983 (500 nM) and additionally, utilising a Halo-Tag-βarr2 construct lacking the finger loop region (dFLR). Angiotensin II (AngII)-induced dynamic BRET changes are shown as concentration-response curves. To emphasise the impact of PKC on βarr2 recruitment, dynamic BRET changes of **Figure 4 a-e** in absence **(b)** or presence of the PKC inhibitor Gö6983 (500 nM) **(c)** are shown as concentration-response curves. Downward arrows and dashed lines indicate the loss in dynamic recruitment by PKC inhibition. **d-h** Corresponding to the data shown in **Figure 4 d and e**, ΔQ-GRK cells were transfected with Angiotensin 1 receptor (AT1R)-Nano luciferase (NanoLuc) and Halo-Tag-β-arrestin2 (βarr2). The concentration-dependent recruitment of βarr2 was measured in the presence of co-transfected GRK5 and 6 kinase dead (KD, K215R) mutants **(d, f)** or in ΔGRK2/3/6 or ΔGRK2/3/5 **(e, g)**. Recruitment data generated for Control + EV, ΔQ-GRK + EV, and ΔQ-GRK + GRK5 or 6 overexpression **(Figure 4 a, b, d, e)** are depicted again to allow comparability. All data points are calculated as Δ net BRET fold change normalised to baseline values and vehicle control, represented as the mean of at least three independent measurements ± SEM. The data are also presented in respective bar graphs, displaying the BRET ratios measured before (baseline) and after stimulation with 1 μM AngII (stimulated), normalised to the basal BRET ratio derived from ΔQ-GRK + EV **(e, g)**. **h** Representative western blot showing the overexpression of GRK5 or 6 and GRK5- or 6-KD mutants in ΔQ-GRK.

**Supplementary Figure 8:**
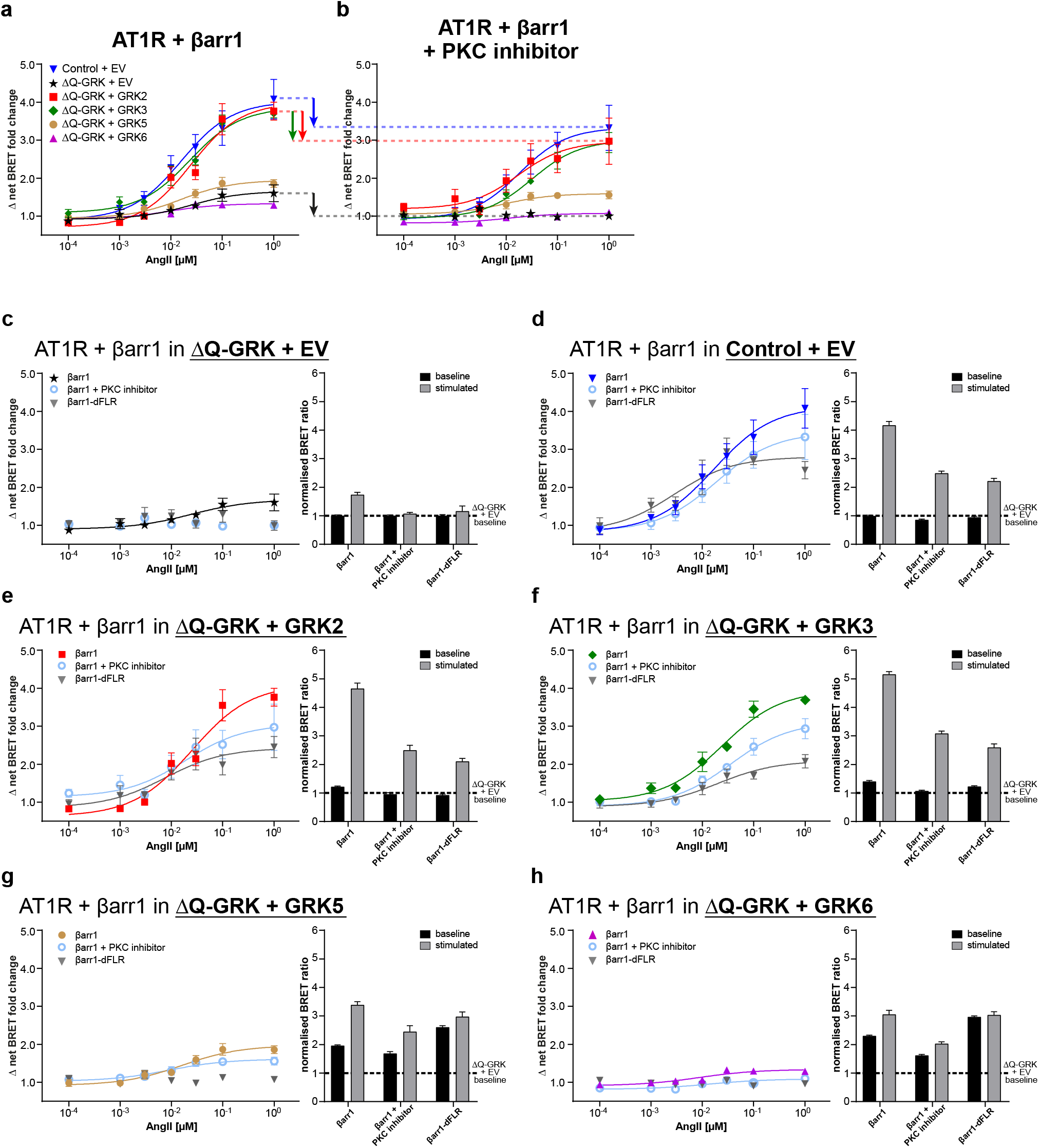
**a, b** ΔQ-GRK or Control cells were transfected with Angiotensin 1 receptor (AT1R)-Nano luciferase (NanoLuc), Halo-Tag-β-arrestin1 (βarr1) and either GRK2, 3, 5, 6, or the empty vector (EV) as indicated. Upon stimulation with Angiotensin II (AngII), dynamic BRET changes are shown as concentration-response curves recorded in absence **(a)** or presence of the PKC inhibitor Gö6983 (500 nM) **(b)**. Downward arrows and dashed lines indicate the loss in dynamic recruitment by PKC inhibition. **c-h** To emphasise the contribution of the individual GRKs to the complex formation of βarr1 with AT1R, data shown in **(a, b)** and **Supplementary Figure 3g** left panel, are now divided into smaller panels according to the individual GRK condition. Additionally, an analogous measurement using a Halo-Tag-βarr1 construct lacking the finger loop region (dFLR) was performed for each condition to test for the formation of a “hanging” complex. All data points are calculated as Δ net BRET fold change normalised to baseline values and vehicle control, represented as the mean of at least three independent measurements ± SEM. Data of these panels are also presented in respective bar graphs, displaying the BRET ratios measured before (baseline) and after stimulation with 1 μM AngII (stimulated), normalised to the basal BRET ratio derived from the corresponding ΔQ-GRK + EV condition.

**Supplementary Figure 9:**
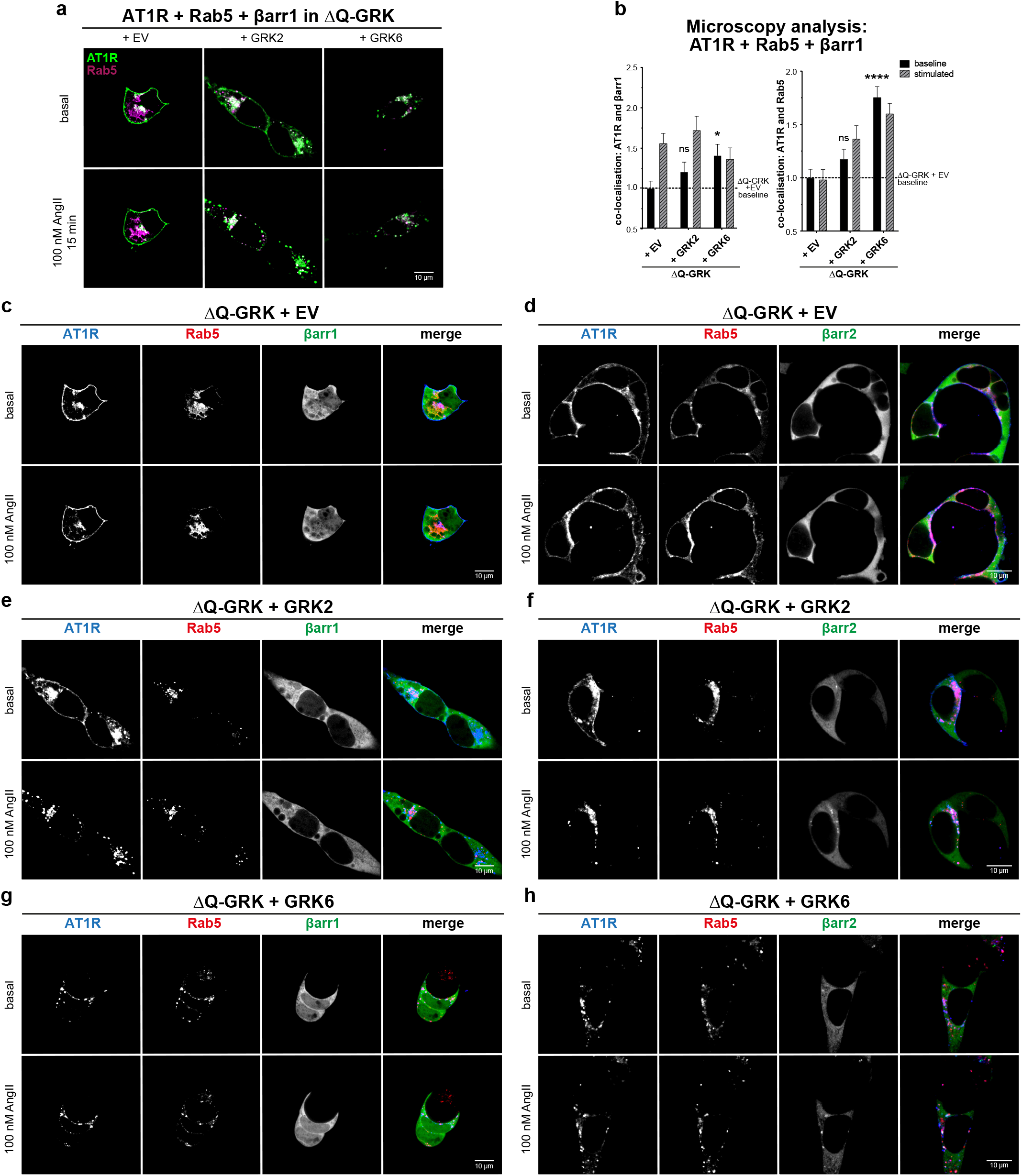
Corresponding to **Figure 4g, h**, GRK2- and 6-dependent Angiotensin 1 receptor (AT1R) internalisation in ΔQ-GRK is shown. **a, b** Analogous experiments were performed as depicted in **Figure 4g, h**, but using β-arrestin1 (βarr1). In short: ΔQ-GRK cells were transfected with hsp-AT1R-CFP (green), Rab5-mCherry (magenta), βarr1-YFP, and either empty vector (EV), GRK2, or GRK6. At least 30 images per condition were taken before (basal) and after 15 minutes of 100 nM Angiotensin II (AngII) stimulation. Representative images are shown in **(a)**. The co-localisation of AT1R and βarr1 or Rab5 was quantified using Squassh and Squassh Analyst. Data are presented as mean fold change in co-localisation signal + SEM, normalised to unstimulated ΔQ-GRK + EV condition **(b**, dashed line). Co-localisation under unstimulated conditions was compared using ANVOA and two-sided Dunnett’s test (*p < 0.05; **p < 0.01; ***p < 0.001; ****p < 0.0001; ns (not significant)). **c-h** Depiction of single channels and overlay (merge) of hsp-AT1R-CFP (blue), Rab5-mCherry (red), and βarr1 **(c, e, g)** and 2 **(d, f, h)**-YFP (green). Additionally, cells were transfected with EV **(c, d)**, GRK2 **(e, f)**, or GRK6 **(g, h)**. Representative images of at least 30 images per condition before (basal) and after 15 minutes of 100 nM AngII stimulation are shown.

**Supplementary Figure 10:**
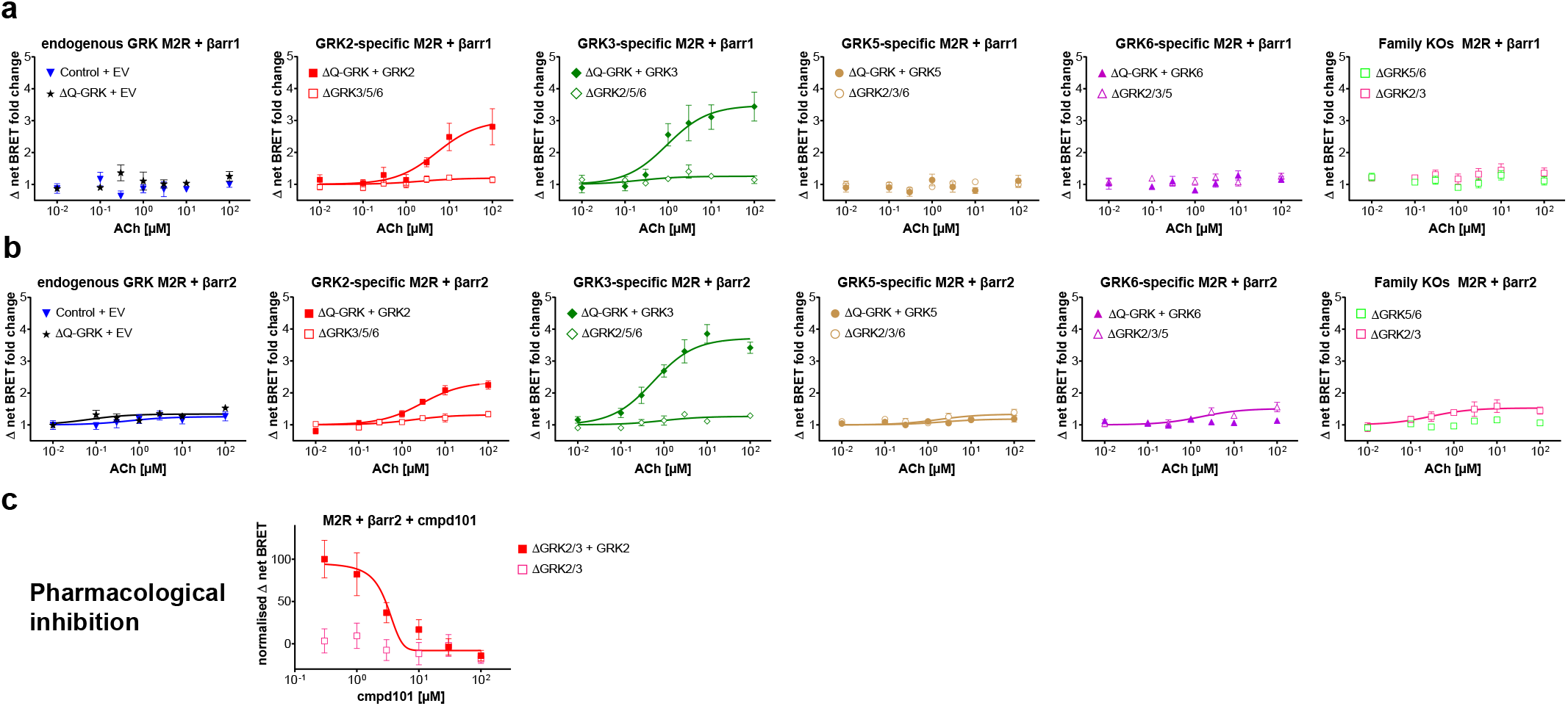
**a, b** The recruitment of β-arrestin (βarr)1 **(a)** or βarr2 **(b)** to the muscarinic M2 acetylcholine receptor (M2R) was measured in presence of no GRK (ΔQ-GRK + empty vector (EV)), all endogenous GRKs (Control + EV), single overexpressed GRKs (ΔQ-GRK + GRK, data identical to **Supplementary Figure 3i**), with only one endogenous GRK left (ΔGRK3/5/6, ΔGRK2/5/6, ΔGRK2/3/6, ΔGRK2/3/5) or with a knockout of the GRK families (ΔGRK2/3 or ΔGRK5/6). Data are presented as Δ net BRET fold change (means of three independent experiments ± SEM). **c** In addition, the experiment using βarr2 was carried out in ΔGRK2/3 cells with or without overexpression of GRK2 that had been pre-incubated with different concentrations of cmpd101 for 10 minutes prior to stimulation with 100 μM of acetylcholine (ACh). Data are presented as normalised Δ net BRET as means of three independent experiments ± SEM.

**Supplementary Figure 11:**
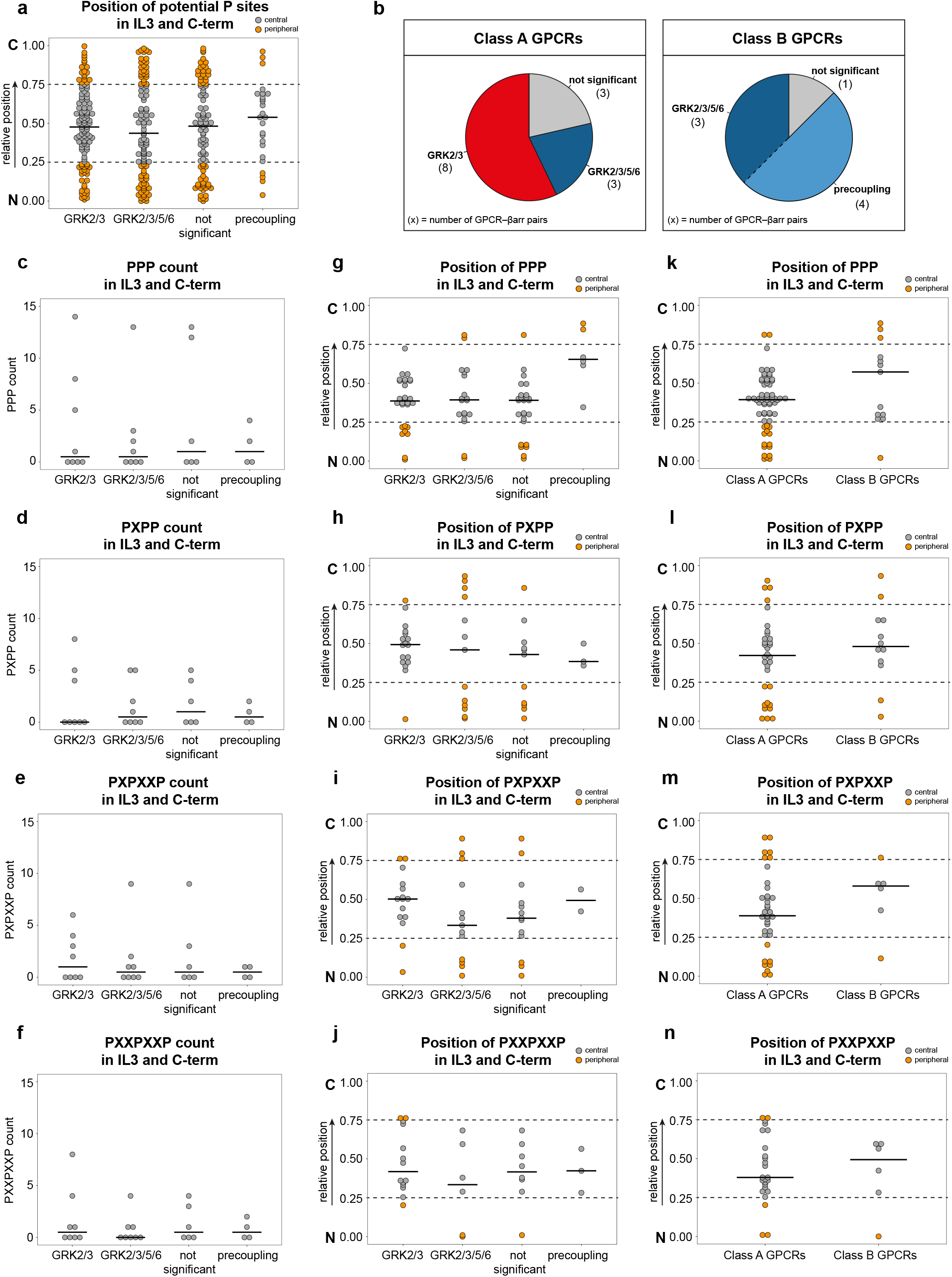
Analysis of the abundance and position of identified putative serine and threonine (Ser/Thr) phosphorylation sites, clusters (PPP, PXPP) and patterns (PXXPXXP, PXPXXP) in the intracellular loop 3 (IL3) and C-terminus (C-term; information from GPCRdb.org) of GPCRs listed in **Supplementary Table 1** with the exception of the unphysiological β2V2 chimera receptor. While X represents any amino acid, P may be a Ser, Thr or a negatively charged amino acid (glutamic acid, Glu; aspartic acid, Asp). Number and positions of potential phosphorylation sites, clusters and patterns were detected using Python 3.8.7. In order to compare positions of potential phosphorylation sites or motifs between GPCRs with varying lengths of IL3 and C-term, their relative position was calculated as the position index in relation to the full length of the respective peptide stretch. Consequently, the relative position of 0 corresponds to the beginning (N), whereas a relative position of 1 corresponds to the end (C) of the respective peptide stretch. All relative positions are displayed as dotplots with corresponding median, indicated by bars. Positions between 0.25 and 0.75 were categorised as central (grey) while positions between 0.0 and 0.25 as well as 0.75 and 1.00 were considered peripheral (orange). Dashed lines mark 0.25 and 0.75 breakpoints. **a** Relative positions of potential phosphorylation sites (Ser/Thr) in the IL3 and C-term of analysed GPCRs grouped according to their GRK-specific β-arrestin (βarr) recruitment. In addition, to the three groups defined in **Figure 2i and j** (GRK2/3-regulated, GRK2/3/5/6-regulated and not significant), GPCRs displaying βarr precoupling (AT1R and V2R) were compiled in a fourth group (precoupling). **b** The analysed receptors were grouped as class A or B according to Oakley *et al.^52^* **(Supplementary Table 1)** and each class is represented as one pie chart. In each pie chart, all GPCR–βarr pairs are assigned to the different GRK-specific βarr recruitment groups. The absolute number of GPCR–βarr pairs presented in each group is indicated in brackets. Since GPCRs displaying βarr precoupling can be considered a subgroup of GRK2/3/5/6-regulated GPCRs, both slices are separated by a dashed line (right pie). **c-f** Abundance of PPP clusters **(c)**, PXPP clusters **(d)**, PXPXXP patterns **(e)** and PXXPXXP patterns **(f)** are displayed as dotplots grouped according to their GRK-specific βarr recruitment. **g-n** Relative positions of P-patterns as indicated. They were grouped according to their GRK-specific βarr recruitment (**g-j**) as well as the GPCR classification (**k-n**).

**Supplementary Table 1:**
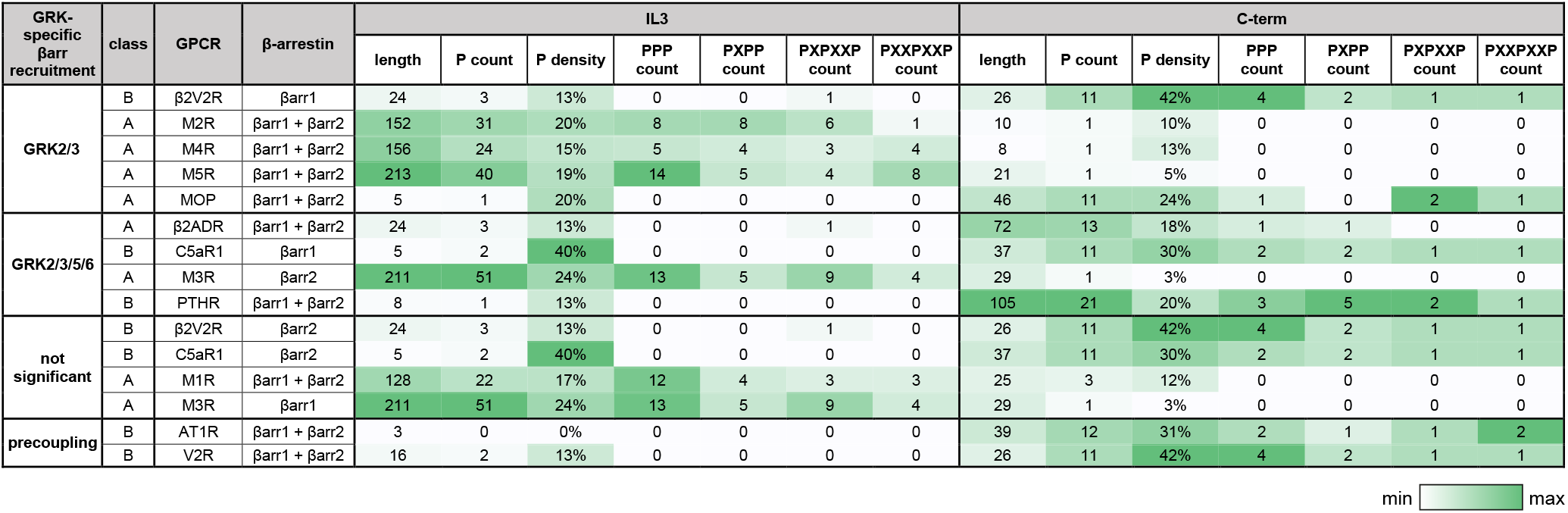
Overview of the length (number of amino acids) of intracellular loop 3 (IL3), C-terminus (C-term) and class (according to Oakley *et al.^52^*) for each analysed GPCR (information from GPCRdb.org) with respective numbers of putative serine and threonine (S/T) phosphorylation sites, clusters (PPP, PXPP) and patterns (PXXPXXP, PXPXXP). While X represents any amino acid, P may be a serine, threonine or a negatively charged amino acid (glutamic acid or aspartic acid). Phosphorylation sites and motifs were detected using Python 3.8.7.

| GRK-specific<br>βarr recruitment | class | GPCR | β-arrestin | IL3 |  |  |  |  |  |  | C-term |  |  |  |  |  |  |
| --- | --- | --- | --- | --- | --- | --- | --- | --- | --- | --- | --- | --- | --- | --- | --- | --- | --- |
|  |  |  |  | length | P count | P density | PPP count | PXPP count | PXPXXP count | PXXPXXP count | length | P count | P density | PPP count | PXPP count | PXPXXP count | PXXPXXP count |
| GRK2/3 | B | β2V2R | βarr1 | 24 | 3 | 13% | 0 | 0 | 1 | 0 | 26 | 11 | 42% | 4 | 2 | 1 | 1 |
|  | A | M2R | βarr1 + βarr2 | 152 | 31 | 20% | 8 | 8 | 6 | 1 | 10 | 1 | 10% | 0 | 0 | 0 | 0 |
|  | A | M4R | βarr1 + βarr2 | 156 | 24 | 15% | 5 | 4 | 3 | 4 | 8 | 1 | 13% | 0 | 0 | 0 | 0 |
|  | A | M5R | βarr1 + βarr2 | 213 | 40 | 19% | 14 | 5 | 4 | 8 | 21 | 1 | 5% | 0 | 0 | 0 | 0 |
|  | A | MOP | βarr1 + βarr2 | 5 | 1 | 20% | 0 | 0 | 0 | 0 | 46 | 11 | 24% | 1 | 0 | 2 | 1 |
| GRK2/3/5/6 | A | β2ADR | βarr1 + βarr2 | 24 | 3 | 13% | 0 | 0 | 1 | 0 | 72 | 13 | 18% | 1 | 1 | 0 | 0 |
|  | B | C5aR1 | βarr1 | 5 | 2 | 40% | 0 | 0 | 0 | 0 | 37 | 11 | 30% | 2 | 2 | 1 | 1 |
|  | A | M3R | βarr2 | 211 | 51 | 24% | 13 | 5 | 9 | 4 | 29 | 1 | 3% | 0 | 0 | 0 | 0 |
|  | B | PTHR | βarr1 + βarr2 | 8 | 1 | 13% | 0 | 0 | 0 | 0 | 105 | 21 | 20% | 3 | 5 | 2 | 1 |
| not significant | B | β2V2R | βarr2 | 24 | 3 | 13% | 0 | 0 | 1 | 0 | 26 | 11 | 42% | 4 | 2 | 1 | 1 |
|  | B | C5aR1 | βarr2 | 5 | 2 | 40% | 0 | 0 | 0 | 0 | 37 | 11 | 30% | 2 | 2 | 1 | 1 |
|  | A | M1R | βarr1 + βarr2 | 128 | 22 | 17% | 12 | 4 | 3 | 3 | 25 | 3 | 12% | 0 | 0 | 0 | 0 |
|  | A | M3R | βarr1 | 211 | 51 | 24% | 13 | 5 | 9 | 4 | 29 | 1 | 3% | 0 | 0 | 0 | 0 |
| precoupling | B | AT1R | βarr1 + βarr2 | 3 | 0 | 0% | 0 | 0 | 0 | 0 | 39 | 12 | 31% | 2 | 1 | 1 | 2 |
|  | B | V2R | βarr1 + βarr2 | 16 | 2 | 13% | 0 | 0 | 0 | 0 | 26 | 11 | 42% | 4 | 2 | 1 | 1 |

min max

**Supplementary Table 2:** Multiple comparisons of GRK-specific β-arrestin recruitment. The p values presented in this table were used to generate the heatmap in **Figure 2 i**. BRET fold changes at saturating ligand concentrations of at least three independent experiments were compared using ANOVA and Bonferroni’s test (* p < 0.05; ** p < 0.01; *** p < 0.001).

| GPCR | βarr | ΔQ-GRK vs. GRK2 |  | ΔQ-GRK vs. GRK3 |  | ΔQ-GRK vs. GRK5 |  | ΔQ-GRK vs. GRK6 |  | ΔQ-GRK vs. Control |  |
| --- | --- | --- | --- | --- | --- | --- | --- | --- | --- | --- | --- |
|  |  | non adj. | adj. | non adj. | adj. | non adj. | adj. | non adj. | adj. | non adj. | adj. |
| AT1R | βarr1 | 0,004 | 0,040 * | 0,006 | 0,050 | 0,687 | 1,000 | 0,618 | 1,000 | 0,001 | 0,008 ** |
| AT1R | βarr2 | 0,155 | 1,000 | 0,117 | 1,000 | 0,000 | 0,000 *** | 0,000 | 0,000 *** | 0,002 | 0,015 * |
| β2ADR | βarr1 | 0,024 | 0,220 | 0,005 | 0,042 * | 0,011 | 0,098 | 0,026 | 0,230 | 0,010 | 0,088 |
| β2ADR | βarr2 | 0,003 | 0,031 * | 0,004 | 0,034 * | 0,017 | 0,155 | 0,009 | 0,079 | 0,086 | 0,770 |
| β2V2R | βarr1 | 0,000 | 0,002 ** | 0,000 | 0,001 *** | 0,089 | 0,801 | 0,619 | 1,000 | 0,078 | 0,704 |
| β2V2R | βarr2 | 0,064 | 0,573 | 0,064 | 0,572 | 0,271 | 1,000 | 0,570 | 1,000 | 0,216 | 1,000 |
| C5aR1 | βarr1 | 0,000 | 0,001 ** | 0,000 | 0,000 *** | 0,000 | 0,002 ** | 0,007 | 0,064 | 0,000 | 0,000 *** |
| C5aR1 | βarr2 | 0,180 | 1,000 | 0,318 | 1,000 | 0,367 | 1,000 | 0,870 | 1,000 | 0,063 | 0,566 |
| M1R | βarr1 | 0,079 | 0,713 | 0,149 | 1,000 | 0,001 | 0,013 * | 0,175 | 1,000 | 0,360 | 1,000 |
| M1R | βarr2 | 0,073 | 0,658 | 0,399 | 1,000 | 0,368 | 1,000 | 0,030 | 0,273 | 0,081 | 0,731 |
| M2R | βarr1 | 0,030 | 0,274 | 0,005 | 0,043 * | 0,827 | 1,000 | 0,882 | 1,000 | 0,700 | 1,000 |
| M2R | βarr2 | 0,012 | 0,104 | 0,000 | 0,000 *** | 0,182 | 1,000 | 0,126 | 1,000 | 0,300 | 1,000 |
| M3R | βarr1 | 0,050 | 0,447 | 0,825 | 1,000 | 0,561 | 1,000 | 0,209 | 1,000 | 0,641 | 1,000 |
| M3R | βarr2 | 0,000 | 0,004 ** | 0,001 | 0,007 ** | 0,167 | 1,000 | 0,008 | 0,068 | 0,001 | 0,005 ** |
| M4R | βarr1 | 0,092 | 0,828 | 0,002 | 0,014 * | 0,653 | 1,000 | 0,958 | 1,000 | 0,138 | 1,000 |
| M4R | βarr2 | 0,007 | 0,066 | 0,000 | 0,002 ** | 0,818 | 1,000 | 0,650 | 1,000 | 0,873 | 1,000 |
| M5R | βarr1 | 0,001 | 0,009 ** | 0,000 | 0,000 *** | 0,977 | 1,000 | 0,929 | 1,000 | 0,000 | 0,000 *** |
| M5R | βarr2 | 0,000 | 0,000 *** | 0,000 | 0,000 *** | 0,903 | 1,000 | 0,710 | 1,000 | 0,000 | 0,004 ** |
| MOP | βarr1 | 0,079 | 0,707 | 0,141 | 1,000 | 0,633 | 1,000 | 0,456 | 1,000 | 0,169 | 1,000 |
| MOP | βarr2 | 0,017 | 0,152 | 0,004 | 0,035 * | 0,540 | 1,000 | 0,922 | 1,000 | 0,510 | 1,000 |
| PTHR | βarr1 | 0,009 | 0,082 | 0,000 | 0,002 ** | 0,015 | 0,134 | 0,004 | 0,039 * | 0,001 | 0,008 ** |
| PTHR | βarr2 | 0,001 | 0,007 ** | 0,002 | 0,016 * | 0,005 | 0,046 * | 0,005 | 0,048 * | 0,000 | 0,000 *** |
| V2R | βarr1 | 0,894 | 1,000 | 0,480 | 1,000 | 0,740 | 1,000 | 0,029 | 0,260 | 0,000 | 0,001 *** |
| V2R | βarr2 | 0,236 | 1,000 | 0,125 | 1,000 | 0,105 | 0,946 | 0,031 | 0,278 | 0,005 | 0,044 * |

| GPCR | βarr | Control vs. GRK2 |  | Control vs. GRK3 |  | Control vs. GRK5 |  | Control vs. GRK6 |  |
| --- | --- | --- | --- | --- | --- | --- | --- | --- | --- |
|  |  | non adj. | adj. | non adj. | adj. | non adj. | adj. | non adj. | adj. |
| AT1R | βarr1 | 0,400 | 1,000 | 0,340 | 1,000 | 0,002 | 0,017 * | 0,000 | 0,003 ** |
| AT1R | βarr2 | 0,326 | 1,000 | 0,406 | 1,000 | 0,000 | 0,000 *** | 0,000 | 0,000 *** |
| β2ADR | βarr1 | 0,448 | 1,000 | 0,966 | 1,000 | 0,709 | 1,000 | 0,435 | 1,000 |
| β2ADR | βarr2 | 0,105 | 0,948 | 0,115 | 1,000 | 0,392 | 1,000 | 0,235 | 1,000 |
| β2V2R | βarr1 | 0,006 | 0,053 | 0,002 | 0,022 * | 0,941 | 1,000 | 0,183 | 1,000 |
| β2V2R | βarr2 | 0,476 | 1,000 | 0,475 | 1,000 | 0,881 | 1,000 | 0,083 | 0,747 |
| C5aR1 | βarr1 | 0,243 | 1,000 | 0,963 | 1,000 | 0,145 | 1,000 | 0,004 | 0,035 * |
| C5aR1 | βarr2 | 0,543 | 1,000 | 0,333 | 1,000 | 0,288 | 1,000 | 0,047 | 0,420 |
| M1R | βarr1 | 0,442 | 1,000 | 0,652 | 1,000 | 0,020 | 0,179 | 0,717 | 1,000 |
| M1R | βarr2 | 0,953 | 1,000 | 0,324 | 1,000 | 0,352 | 1,000 | 0,591 | 1,000 |
| M2R | βarr1 | 0,015 | 0,132 | 0,002 | 0,021 * | 0,867 | 1,000 | 0,812 | 1,000 |
| M2R | βarr2 | 0,002 | 0,014 * | 0,000 | 0,000 *** | 0,745 | 1,000 | 0,586 | 1,000 |
| M3R | βarr1 | 0,120 | 1,000 | 0,806 | 1,000 | 0,301 | 1,000 | 0,092 | 0,832 |
| M3R | βarr2 | 0,899 | 1,000 | 0,829 | 1,000 | 0,008 | 0,071 | 0,172 | 1,000 |
| M4R | βarr1 | 0,004 | 0,035 * | 0,000 | 0,001 *** | 0,338 | 1,000 | 0,151 | 1,000 |
| M4R | βarr2 | 0,010 | 0,089 | 0,000 | 0,002 ** | 0,943 | 1,000 | 0,768 | 1,000 |
| M5R | βarr1 | 0,041 | 0,366 | 0,349 | 1,000 | 0,000 | 0,000 *** | 0,000 | 0,000 *** |
| M5R | βarr2 | 0,187 | 1,000 | 0,086 | 0,772 | 0,000 | 0,004 ** | 0,001 | 0,009 ** |
| MOP | βarr1 | 0,005 | 0,049 * | 0,010 | 0,093 | 0,350 | 1,000 | 0,502 | 1,000 |
| MOP | βarr2 | 0,058 | 0,525 | 0,014 | 0,122 | 0,962 | 1,000 | 0,573 | 1,000 |
| PTHR | βarr1 | 0,222 | 1,000 | 0,472 | 1,000 | 0,147 | 1,000 | 0,396 | 1,000 |
| PTHR | βarr2 | 0,039 | 0,351 | 0,017 | 0,149 | 0,006 | 0,052 | 0,006 | 0,050 * |
| V2R | βarr1 | 0,000 | 0,001 *** | 0,000 | 0,002 ** | 0,000 | 0,000 *** | 0,000 | 0,000 *** |
| V2R | βarr2 | 0,001 | 0,005 ** | 0,000 | 0,002 ** | 0,000 | 0,002 ** | 0,000 | 0,001 *** |

**Supplementary Table 3:** Complete results of statistical analysis of BRET ratios shown in **Figure 4** and **Supplementary Figure 7 a, 8 c-h**. Statistical significance was determined by two-way mixed model ANOVA followed by paired t-test comparing stimulated BRET ratios to their corresponding baseline as well as Tukey’s test comparing baselines or stimulated BRET ratios between each other (*p < 0.05; **p < 0.01; ***p < 0.001; ****p < 0.0001).

| comparing baseline and stimulated |  |  |  |  |
| --- | --- | --- | --- | --- |
| condition | β-arrestin | + βarr | + βarr-dFLR | + βarr<br>+ PKC inhibitor |
| Control + EV | βarr1 | 0.00444 ** | 0.00829 ** | 0.00300 ** |
| ΔQ-GRK + EV | βarr1 | 0.00469 ** | 0.36109 | 0.57302 |
| ΔQ-GRK + GRK2 | βarr1 | 0.00791 ** | 0.00185 ** | 0.00744 ** |
| ΔQ-GRK + GRK3 | βarr1 | 0.00171 ** | 0.01415 * | 0.00712 ** |
| ΔQ-GRK + GRK5 | βarr1 | 0.00395 ** | 0.00407 ** | 0.01342 * |
| ΔQ-GRK + GRK6 | βarr1 | 0.01012 * | 0.45936 | 0.00920 ** |
| Control + EV | βarr2 | 0.00859 ** | 0.09450 | 0.00030 *** |
| ΔQ-GRK + EV | βarr2 | 0.00033 *** | 0.86111 | 0.00981 ** |
| ΔQ-GRK + GRK2 | βarr2 | 0.00469 ** | 0.05126 | 0.00141 ** |
| ΔQ-GRK + GRK3 | βarr2 | 0.00601 ** | 0.05231 | 0.00347 ** |
| ΔQ-GRK + GRK5 | βarr2 | 0.04058 * | 0.01303 * | 0.02003 * |
| ΔQ-GRK + GRK6 | βarr2 | 0.01930 * | 0.34421 | 0.10205 |

| comparing baseline and baseline |  |  |  |  |
| --- | --- | --- | --- | --- |
| condition | β-arrestin | βarr vs.<br>βarr-dFLR | βarr vs.<br>βarr + PKC<br>inhibitor | βarr-dFLR vs.<br>βarr + PKC<br>inhibitor |
| Control + EV | βarr1 | 0.93184 | 0.62262 | 0.82231 |
| ΔQ-GRK + EV | βarr1 | 1.00000 | 1.00000 | 1.00000 |
| ΔQ-GRK + GRK2 | βarr1 | 0.45148 | 0.51739 | 0.99077 |
| ΔQ-GRK + GRK3 | βarr1 | 0.50462 | 0.14492 | 0.57972 |
| ΔQ-GRK + GRK5 | βarr1 | 0.18168 | 0.69820 | 0.06282 |
| ΔQ-GRK + GRK6 | βarr1 | 0.03082 * | 0.02745 * | 0.00092 *** |
| Control + EV | βarr2 | 0.23070 | 0.50580 | 0.78438 |
| ΔQ-GRK + EV | βarr2 | 1.00000 | 1.00000 | 1.00000 |
| ΔQ-GRK + GRK2 | βarr2 | 0.01233 * | 0.00609 ** | 0.76853 |
| ΔQ-GRK + GRK3 | βarr2 | 0.00600 ** | 0.00945 ** | 0.92324 |
| ΔQ-GRK + GRK5 | βarr2 | 0.00269 ** | 0.00002 **** | 0.00000 **** |
| ΔQ-GRK + GRK6 | βarr2 | 0.11392 | 0.05662 | 0.84639 |

| comparing stimulated and stimulated |  |  |  |  |
| --- | --- | --- | --- | --- |
| condition | β-arrestin | βarr vs.<br>βarr-dFLR | βarr vs.<br>βarr + PKC<br>inhibitor | βarr-dFLR vs.<br>βarr + PKC<br>inhibitor |
| Control + EV | βarr1 | 0.00003 **** | 0.00002 *** | 0.32708 |
| ΔQ-GRK + EV | βarr1 | 0.17696 | 0.11616 | 0.94195 |
| ΔQ-GRK + GRK2 | βarr1 | 0.00051 *** | 0.00014 ** | 0.50894 |
| ΔQ-GRK + GRK3 | βarr1 | 0.00006 **** | 0.00004 *** | 0.18578 |
| ΔQ-GRK + GRK5 | βarr1 | 0.56456 | 0.01106 | 0.41309 |
| ΔQ-GRK + GRK6 | βarr1 | 0.99380 | 0.00593 ** | 0.00694 ** |
| Control + EV | βarr2 | 0.03493 * | 0.12697 | 0.58481 |
| ΔQ-GRK + EV | βarr2 | 0.10554 | 0.33143 | 0.65260 |
| ΔQ-GRK + GRK2 | βarr2 | 0.00629 ** | 0.00779 ** | 0.97985 |
| ΔQ-GRK + GRK3 | βarr2 | 0.01525 * | 0.01205 * | 0.98205 |
| ΔQ-GRK + GRK5 | βarr2 | 0.09398 | 0.00124 ** | 0.01643 * |
| ΔQ-GRK + GRK6 | βarr2 | 0.00648 ** | 0.01708 * | 0.67218 |

## Methods

### Cell culture

HEK293 cells were originally obtained from DSMZ Germany (ACC 305) and cultured in Dulbecco’s modified Eagle’s medium (DMEM; Sigma-Aldrich D6429), complemented with 10 % fetal calf serum (FCS; Sigma-Aldrich F7524) and 1 % of penicillin and streptomycin mixture (Sigma-Aldrich P0781) at 37 °C with 5 % CO_2_. The cells were passaged every 3-4 days. Cells were regularly checked for mycoplasma infections using the LONZA MycoAlert mycoplasma detection kit (LT07-318).

### CRISPR/Cas 9 mediated knockout of GRK2, 3, 5, and 6

Stable GRK knockout cells were generated by transient transfection using self-made PEI reagent (Sigma-Aldrich, 408727, diluted to 10 μg/ml, pH 7.2, adjusted with HCl) of the parental cells (HEK293) with lentiCRISPR v2 plasmid^57^ (Addgene #52961) containing target specific gRNAs listed in **Methods Table 1.** Complementary forward and reverse oligos were annealed and ligated into the BsmBI-restricted lentiCRISPR v2 vector. This vector could also be used for the generation of viral particles, but in our approach, they were directly transfected into the target cells. In order to prevent side effects caused by multiple transfection and selection rounds, all cell clones were created in singular attempts. To knockout one specific GRK, four different gRNA constructs were simultaneously transfected. In line, double, triple, or quadruple knockout cells were generated by transfection of 8, 12, or 16 respective gRNA constructs at once. The transfected cells were then selected using 1 μg/ml puromycin (Sigma-Aldrich #P8833). Limited dilution was used to establish single cell clones, which were then analysed for absence of the target protein by Western Blot analysis. A puromycin selected cell pool transfected with empty lentiCRISPR v2 plasmid was used as Control.

**Methods Table 1:**
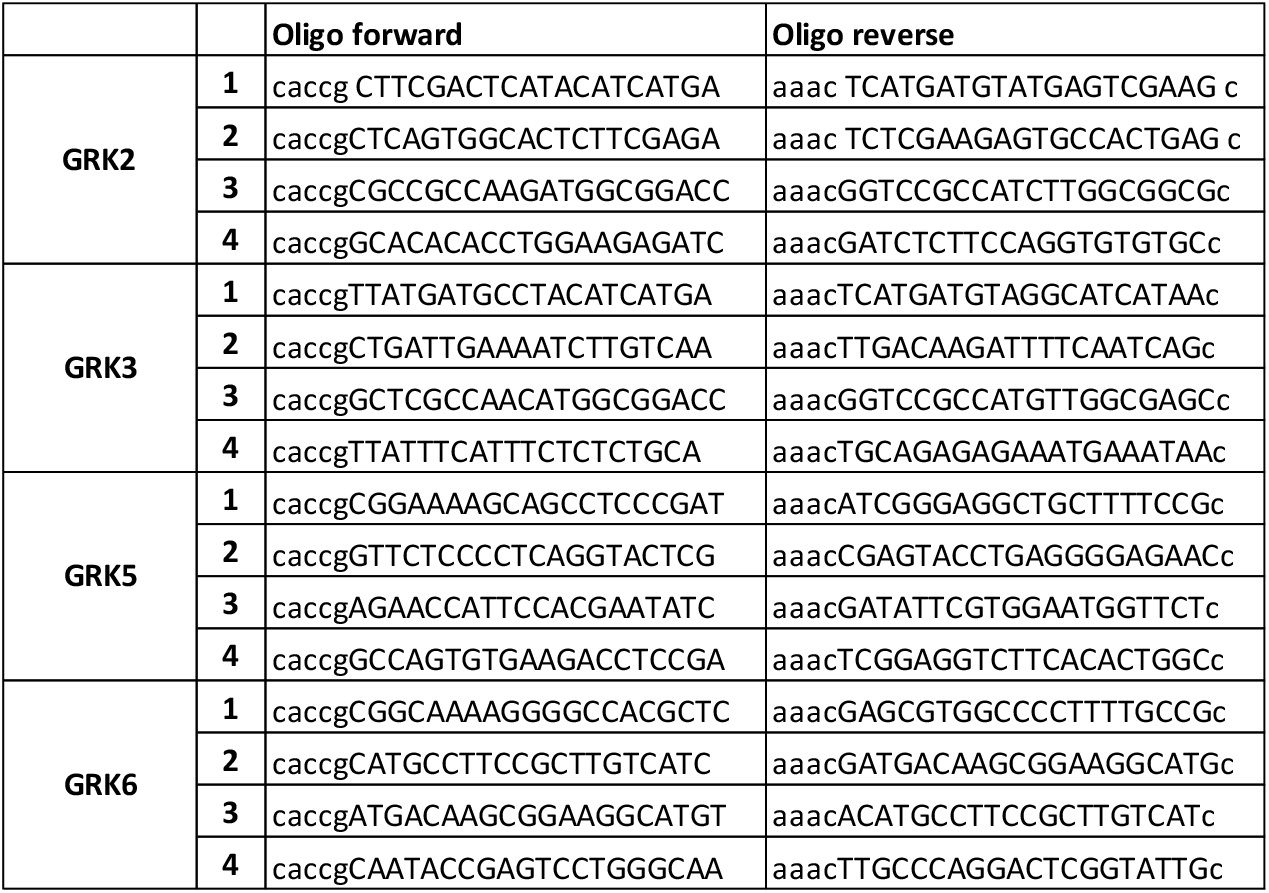
gRNA sequences for targeting different GRKs. gRNA sequence in capital letters, overhangs for ligation in lowercase letters. Sequences of gRNAs were validated in Sanjana *et al*.^57^

### Establishment of GRK expression constructs

The used pcDNA3-GRK2 expression construct was described before^58^. GRK3 (NCBI reference sequence NM_005160.4), GRK5 (NCBI reference sequence NM_005308.3), and GRK6 (NCBI reference sequence NM_001004106.3) were amplified by PCR using GRK-specific primers including restriction sites for HindIII (forward primer) and BamHI (reverse primer) (GRK3: Forward primer (fw) – CTT AAG CTT GCC ACC ATG GCG GAC CTG GAG GCTG, Reverse Primer (rev) – CTT AGG ATC CTA GAG GCC GTT GCT GTT TCTG; GRK5: fw – CTT AAG CTT GCC ACC ATG GAG CTG GAA AAC ATC GTG, rev – CTT AGG ATC CTA GCT GCT TCC GGT GGAG; GRK6: fw – CTT AAG CTT GCC ACC ATG GAG CTC GAG AAC ATC GTAG, rev – CTT AGG ATC CTA GAG GCG GGT GGG GAGC). GRK3 and 6 were amplified from human leucocyte cDNA, GRK5 was amplified from a beta galactosidase fusion plasmid described before^59^. The fragments were ligated into pcDNA3 plasmids after BamHI and HindIII digest. Sequences of all plasmids were validated by sequencing.

The K215R kinase dead (KD) mutants of GRK5 and 6 were created by site-directed mutagenesis.

### Western blot

Cells were washed once with ice-cold PBS and subsequently lysed with RIPA Buffer (1 % NP-40, 1 mM EDTA, 50 mM Tris pH 7.4, 150 mM NaCl, 0.25 % sodium deoxycholate), supplemented with protease and phosphatase inhibitor cocktails (Roche, #04693132001, #04906845001). Cleared lysates were boiled with SDS loading buffer and 15 μg of total protein were loaded onto each lane of 10 % polyacrylamide gels. After transfer onto nitrocellulose membranes, the total protein was detected by using specific antibodies (GRK2: Santa Cruz sc-13143; GRK3: Cell signaling technology #80362; GRK5: Santa Cruz, sc-518005; GRK6: Cell signaling technology #5878, Actin: Sigma-Aldrich A5441). For experiments using the MOP phospho-specific antibodies see below. Quantification of the blots was done using Fujifilm multi Gauge software (V3.0).

### Cell viability assay and growth rates determination

Cells were seeded with a density of 5,000 cells / 100 μl in 96-well plates, with 12-16 wells per cell line. After 48 hours of incubation at 37 °C and 5 % CO_2_, 20 μl of the cell titer blue reagent (Promega, G8081) were added to 3-4 wells for each cell line, and the cells were further incubated for 1.5 hours under the same growth conditions. After the incubation time, fluorescence (exitation 540 nm, emission 610 nm, Gain 40, top reading) was measured using a TECAN Infinite 200 (Tecan, Crailsheim, Germany) plate reader. The plate was further incubated and 72 hours, 96 hours, and 120 hours after seeding, three wells were measured as just described. The proliferation was reported as relative fluorescence signal compared to the first measurement (48 hours after seeding) ± standard error of the mean (SEM) of at least n=3 experiments. Using these data, the growth rates were determined for the three 24 hours intervals and mean values ± SEM are reported.

### Stable MOP-expressing cells and immunoprecipitation experiments

A retroviral expression vector was created by replacing the mCherry expression cassette of pMSCV-IRES-mCherry FP (a gift from Dario Vignali, Addgene plasmid #52114) with a neomycine resistance cassette of pcDNA3 plasmid resulting in empty pMSCV-IRES-NEO vector. The open reading frame (ORF) of N-terminal haemagglutinin (HA) tagged murine μ-opioid receptor (MOP) gene^59^ was inserted into the multi-cloning-site of this vector resulting in pMSCV-HA-MOP-IRES-NEO. Freshly produced retroviral particles were used to transduce GRK-KO cell clones or respective control cells. The cells were selected with 1 mg/ml G418 (Gibco, 11811-031) for ten days and subsequently stained with Anti-HA-IgG CF-488A (Sigma-Aldrich, SAB4600054) antibody and sorted for equal surface expression of the MOP by flow cytometry.

For immunoprecipitation experiments, 4 × 10^6^ cells were seeded in a 21 cm^2^ dish and after 24 hours stimulated for ten minutes with 10 μM DAMGO ([D-Ala^2^, *N*-MePhe^4^, Gly-ol]-enkephalin, Tocris 1171) in DMEM medium (Sigma-Aldrich, D6429) without supplements. After washing with ice-cold PBS, the cells were lysed in 500 μl RIPA buffer as described in **“Western Blot” section**. 400 μg of total protein lysates were incubated with 20 μl of HA-beads slurry (Thermo Scientific, 26182) at 4 °C on a turning wheel for two hours. The beads were then washed three times with RIPA buffer containing protease and phosphatase inhibitors and 75 μl sample buffer (125 mM Tris pH 6.8, 4 % SDS, 10 % glycerol, 167 mM DTT) were added and the samples were heated to 42 °C for 20 minutes. The samples were loaded onto 10 % polyacrylamide gels (7.5 μl per 15-well mini gel lane) and phosphorylation was detected using self-made phospho-specific antibodies (pS^363^, pT^370^, pT^379^, pS^375^, pT^376^)^20,21^. The total receptors were detected with an anti HA-antibody (Cell signaling technology # 3724). Quantification of the blots was done using Fujifilm multi Gauge software (V3.0).

### Intermolecular bioluminescence resonance energy transfer (BRET)

The GRK-selective β-arrestin recruitment assay was performed either in Control or specific ΔGRK cell lines. In 21 cm^2^ dishes, 1.6 × 10^6^ cells were seeded and transfected the next day with 0.5 μg of the respective GPCR C-terminally fused to Nano luciferase (NanoLuc), 1 μg of β-arrestin constructs N-terminally fused to a Halo-ligand binding Halo-Tag and 0.25 μg of one GRK or empty vector. In case of the PTH1R and V2R, the BRET pair was swapped. All transfections were conducted following the Effectene transfection reagent manual by Qiagen (#301427) and then incubated at 37 °C overnight. Into poly-D-lysine-coated 96-well plates (Brand, 781965), 40,000 cells were seeded per well in presence of Halo-ligand (Promega, G980A) at a ratio of 1:2,000. A mock labelling condition without addition of the Halo-ligand was seeded for each transfection. After 24 hours, the cells were washed twice with measuring buffer (140 mM NaCl, 10 mM HEPES, 5.4 mM KCl, 2 mM CaCl_2_, 1 mM MgCl_2_; pH 7.3) and NanoLuc-substrate furimazine (Promega, N157B) was added in a ratio of 1:35,000 in measuring buffer. A Synergy Neo2 plate reader (Biotek) with a custom-made filter (excitation bandwidth 541-550 nm, emission 560-595 nm, fluorescence filter 620/15 nm) was used to perform the measurements. The baseline was monitored for three minutes. After addition of the respective agonist, the measurements were continued for five minutes. By subtracting the values measured for mock labelling conditions, the initial BRET change was corrected for labelling efficiency. Halo-corrected BRET changes were calculated by the division of the corrected and averaged values measured after ligand stimulation by the respective, corrected and averaged baseline values. Subsequently, this corrected BRET change was divided by the vehicle control for the final dynamic Δ net BRET change.

For the analysis of the respective BRET ratios before and after stimulation, the Halo-corrected and averaged BRET ratios before stimulation (baseline) and the Halo-corrected and averaged BRET ratios after stimulation with saturating ligand concentration (stimulated) are displayed as bar graphs.

Receptors were stimulated as follows: human Angiotensin 1 receptor (AT1R) with Angiotensin II (AngII; Tocris 4474-91-3, in measuring buffer), human β2 adrenergic receptor (β2ADR) with isoproterenol (Iso; Sigma-Aldrich I5627, in water) Epinephrine (Sigma-Aldrich E4642, in measuring buffer) and Norepinephrine (Sigma-Aldrich 74488, in measuring buffer), human β2ADR with an exchanged C-terminus of the vasopressin type 2 receptor (β2V2), with Iso, human complement 5a receptor 1 (C5aR1) with C5aR-agonist (AnaSpec AS65121, in measuring buffer), human muscarinic 1,2,3,4, and 5 acetylcholine receptors (M1R, M2R, M3R, M4R, and M5R) with acetylcholine (ACh; Sigma-Aldrich A6625, in measuring buffer), murine μ-opioid receptor (MOP) with [D-Ala^2^, *N*-MePhe^4^, Gly-ol]-enkephalin (DAMGO; Tocris 1171, in water), human parathyroid hormone 1 receptor (PTH1R) with parathyroid-hormone (1-34) (PTH(1-34); Bachem 4011474, in measuring buffer), and human vasopressin type 2 receptor (V2R) with [Arg^8^]-vasopressin (AVP; Tocris 2935, in water). The transfected β-arrestins are of bovine origin. β-arrestin1 constructs lacking the fingerloop region (dFLR) were generated as described in Cahill *et al.^31^* by site-directed mutagenesis. The corresponding β-arrestin2 constructs were designed homologously.

If not further elaborated, the utilised cDNAs were obtained from cDNA resource center (www.cDNA.org) or Addgene. The Halo-Tag or NanoLuc genes were acquired from Promega and were genetically fused to the respective N- or C-termini.

In case of cmpd101 (Tocris 15777006, in DMSO) and pindolol (Sigma-Aldrich, P0778, in 0.1 M HCl) inhibitor experiments, the β-arrestin recruitment was induced with either 1 μM isoproterenol (in case of the β2ADR) or 10 μM ACh (in case of the M2R) after a ten-minute incubation period with different concentrations of cmpd101 or pindolol.

In case of the Gö6983 (Tocris, 2285, in DMSO) experiments, the cells were pre-incubated with 500 nM of the inhibitor at 37 °C for 1 h, and subsequently stimulated with different concentrations of AngII.

### Fluorometric assessment of GRK expression and functionality of GRK-YFP constructs

For the assessment of GRK expression levels, C-terminal GRK-YFP fusions were constructed via isothermal plasmid assembly, keeping the identical vector backbone. 1.6 × 10^6^ ΔQ-GRK cells were seeded in 21 cm^2^ dishes and transfected the next day with 1.5 μg of either GRK2-, 3-, 5-, or 6-YFP fusion constructs. All transfections were conducted following the Effectene transfection reagent manual by Qiagen (#301427) and then incubated at 37 °C overnight. Into poly-D-lysine-coated 96-well plates (Brand, 781965), 40,000 cells were seeded per well. After 24 hours of incubation at 37°C, the fluorescence was assessed using a Synergy Neo2 plate reader (Biotek) and a corresponding YFP filter (excitation bandwidth 465-505 nm, emission 496-536 nm). To confirm the catalytic activity of the used GRK-YFP fusion constructs, a GRK-specific β-arrestin2 recruitment intermolecular BRET assay featuring the PTH1R-Halo-Tag and β-arrestin2-NanoLuc was performed. In this case, cells were transfected as described in **“intermolecular bioluminescence resonance energy transfer (BRET)” section**. Instead of untagged GRK constructs, 0.25 μg of the GRK-YFP fusion constructs were transfected. After stimulation with PTH(1-34), the data were recorded and processed as described above.

### Statistical analysis of intermolecular BRET

BRET ratios and fold changes are displayed as mean of at least three independent experiments with error bars indicating the SEM. Statistical analysis was performed using analysis of variance (ANOVA; one-way or two-way mixed model ANOVA) as well as appropriate multiple comparisons as indicated in corresponding figure legends. Data were prepared using Python 3.8.7 and statistical analysis was conducted in R 4.0.3. A type I error probability of 0.05 was considered to be significant in all cases. Two-way mixed model ANOVA was performed using *ez* R package (Lawrence, MA. (2011) ez: Easy analysis and visualization of factorial experiments. R package version 4.4-0. http://CRAN.R-project.org/package=ez) and multiple comparisons were conducted using the *multcomp* R package^60^. The clustering heatmap was generated using the *pheatmap* R package (Kolde, R. (2013). pheatmap: Pretty Heatmaps. R package version 1.0.12. http://CRAN.R-project.org/package=pheatmap.).

### Intramolecular BRET

ΔQ-GRK or Control cells were transfected with 1.2 μg untagged β2ADR, 0.12 μg of β-arrestin2 FlAsH5-tagged biosensor C-terminally coupled to NanoLuc, 0.25 μg of either GRK2, 3, 5, 6, or empty vector as noted, following the Effectene transfection reagent protocol by Qiagen. 24 hours after transfection, 40,000 cells were seeded per well into poly-D-lysine coated 96-well plates and incubated overnight at 37 °C. For this study, the FlAsH (fluorescine arsenical hairpin-binder)-labelling procedure previously described Hoffmann *et al.^61^* was adjusted for 96-well plates. Before the FlAsH-labelling procedure, the cells were washed twice with PBS, then incubated with 250 nM FlAsH in labelling buffer (150 mM NaCl, 10 mM HEPES, 25 mM KCl, 4 mM CaCl_2_, 2 mM MgCl_2_, 10 mM glucose; pH7.3), complemented with 12.5 μM 1,2-ethane dithiol (EDT) for sixty minutes at 37 °C. After aspiration of the FlAsH labelling or mock labelling solutions, the cells were incubated for 10 minutes at 37 °C with 100 μl 250 μM EDT in labelling buffer per well. Addition of the NanoLuc substrate, measurement and analysis of the BRET change was performed as described above (see **“intermolecular bioluminescence resonance energy transfer (BRET)” section**).

### Microscopy

The morphology of the generated cell clones was documented during regular cell culture procedure using phase contrast microscopy at the invitrogen EVOS FL Auto in 10 × magnification.

Receptor internalisation of fixed cells stably expressing the MOP was analysed using confocal microscopy. ΔGRK2/3, ΔGRK5/6, ΔQ-GRK, and control cells were grown on poly-L-lysine-coated coverslips for 2-3 days. After the treatment with 10 μM DAMGO at 37 °C for 30 minutes, cells were fixed with 4 % paraformaldehyde and 0.2 % picric acid in phosphate buffer (pH 6.9) for 30 minutes at room temperature. Then coverslips were washed several times with PBS w/o Ca^2+^/Mg^2+^ buffer. After washing with 50 % and 100 % methanol for 3 minutes, cells were permeabilised with phosphate buffer for 2 hours and then incubated with self-made anti-HA antibody (1:500) followed by Alexa488-conjugated secondary antibody (1:2,000) (Life Technologies, Thermo Fisher Scientific A11008). Specimens were mounted with Roti®-Mount FluorCare DAPI (Carl Roth, HP20.1) and examined using a Zeiss LSM510 META laser scanning confocal microscope.

Live cell experiments were performed to record the translocation of PTH1R, M5R, V2R or AT1R, β-arrestin, and the early endosome marker Rab5 upon agonist stimulation at a Leica SP8 laser scanning confocal microscope. Therefore ΔGRK2/3, ΔGRK5/6, ΔQ-GRK, and control cells were transfected with 1 μg of the C-terminally CFP-fused receptor (in the case of AT1R, we included an hsp-export tag^62^ and transfected 1.25 μg), 0.5 μg of β-arrestin-YFP, 0.5 μg of Rab5-mCherry (kindly provided by Tom Kirchhausen (Harvard Medical School, Boston, USA)) and 0.25 μg GRK expression constructs (as indicated) in a 21 cm^2^ dish, according to the Effectene transfection reagent manual by Qiagen. After 24 hours, 700,000 transfected cells were seeded onto poly-D-lysine-coated glass cover slips in 6-well plates. Another 24 hours later, the cover slips were washed twice with measuring buffer and subsequently imaged before and after stimulation for 15 min with either 100 nM PTH(1-34), 100 μM ACh, 100 nM AVP or 100 nM AngII as indicated. CFP was excited at a wavelength of 442 nm, YFP at 514 nm, and mCherry at 561 nm. The images were acquired with a 63 × water immersion objective, with zoom factor 3, line average 3, and 400 Hz in a 1024 × 1024 pixel format. Subcellular features of the acquired images were segmented and quantified using an ImageJ-based software called segmentation and quantification of subcellular shapes (Squassh). Utilising Squassh’s deconvolution, denoising, and segmentation of the three fluorescence channels present in each image, the raw data readout was then eligible for analysis using the R-based software SquasshAnalyst as described by A. Rizk^63,64^. All image-derived data in this study were processed and analysed with this method and are presented as fold change in co-localisation signal. Statistical analysis of quantified microscopy data was performed using GraphPad Prism 7.03. Unstimulated and stimulated co-localisation was compared using paired t-test. To identify significantly increased co-localisation under unstimulated conditions, quantified co-localisation was compared using ANOVA and two-sided Dunnett’s test.

### Enzyme-linked immunosorbent assay (ELISA)

Receptor internalisation was quantified using a linear surface receptor enzyme-linked immunosorbent assay (ELISA) that has been characterised extensively^65,66^. Equal numbers of ΔGRK2/3, ΔGRK5/6, ΔQ-GRK, and control HEK293 cells stably expressing HA-tagged murine MOP were seeded onto poly-L-lysine-coated 24-well plates for 2-3 days. Then, cells were pre-incubated with 1 μg/ml anti-HA antibody for 2 h at 4 °C. After 30 minutes treatment with 10 μM DAMGO at 37 °C, the cells were fixed with 4 % paraformaldehyde and 0.2 % picric acid in phosphate buffer (pH 6.9) for 30 minutes at room temperature and incubated with peroxidase-conjugated anti-rabbit antibody (Cell Signaling technology #7074) overnight at 4 °C. After washing, the plates were developed with ABTS solution (Sigma-Aldrich A3219) and analysed at 405 nm using a microplate reader.

### Label-free Dynamic Mass Redistribution (DMR) biosensing

Dynamic mass redistribution (DMR) experiments were performed using the Corning Epic (Corning, NY, USA) biosensor technology as previously described in detail^67–73^. In short, for DMR detection 9 × 10^5^ ΔQ-GRK cells were seeded into 21 cm^2^ dishes and cultured until reaching a confluence of 60-80%, which is critical for the maintenance of a consistent proliferation phenotype and for comparable transfection efficiencies. Subconfluent cells were then transiently transfected with empty vector (pcDNA3) or expression plasmids (pcDNA3-based) encoding for the AT1R (hsp-AT1R-CFP), β-arrestin2, GRK2 or GRK6. The next day, transfected cells were seeded at a density of 2 × 10^4^ cells per well into Corning Epic biosensor microplates and incubated overnight (37 °C, 5 % CO_2_). Prior to DMR detection, cells were washed three times with HBSS buffer containing 20 mM HEPES and 0.1 % fatty acid-free bovine serum albumin (BSA) (Sigma-Aldrich). Cells were then placed into the Epic-DMR-reader and equilibrated for 1 hour at 37 °C to achieve baseline stabilisation. Compounds diluted in the same buffer were added to the biosensor plate using the CyBio Selma semi-automatic pipettor (Analytik Jena AG) and ligand-induced DMR alterations were monitored as picometer (pm) wavelength shifts for at least 3600 seconds in 15 seconds intervals. Real-time DMR recordings are means + SEM of three technical replicates and were corrected by the pm wavelength shifts obtained in empty vector transfectants. Concentration-effect curves are means ± SEM of three to four independent biological replicates and were derived from the area under the curve (AUC) between zero and 1800 seconds using a three parameter logistic equation and the GraphPad Prism (8.4.3) software. Curves were fitted with ‘bottom’ constrained to zero while all other settings were left to their default values. Two way ANOVA was used for statistical analysis.

## Abbreviations

ACh: acetylcholine
AngII: Angiotensin II
ANOVA: analysis of variance
AT1R: Angiotensin 1 receptor
AU: arbitrary units
AUC: area under the curve
AVP: [Arg^8^]-vasopressin
βarr1: β-arrestin1
βarr2: β-arrestin2
β2ADR: β2 adrenergic receptor
β2V2: β2ADR with an exchanged C-terminus of the vasopressin type 2 receptor
BRET: bioluminescence resonance engergy transfer
C5aR1: complement 5a receptor 1
CFP: cyan fluorescent protein
cmpd101: GRK2/3 inhibitor
DAMGO: [D-Ala2, N-MePhe4, Gly-ol]-enkephalin
ΔGRK2: GRK2 knockout in HEK293
ΔGRK2/3: GRK2 and 3 double knockout in HEK293
ΔGRK2/3/5: GRK2, 3 and 5 triple knockout in HEK293
ΔGRK2/3/6: GRK2, 3 and 6 triple knockout in HEK293
ΔGRK2/5/6: GRK2, 5 and 6 triple knockout in HEK293
ΔGRK3: GRK3 knockout in HEK293
ΔGRK3/5/6: GRK3, 5 and 6 triple knockout in HEK293
ΔGRK5: GRK5 knockout in HEK293
ΔGRK5/6: GRK5 and 6 double knockout in HEK293
ΔGRK6: GRK6 knockout in HEK293
ΔQ-GRK: GRK2, 3, 5, 6 quadruple knockout in HEK293
dFLR: deleted finger loop region
DMEM: Dulbecco’s modified eagle medium
DTT: Dithiothreitol
EDT: 1,2-ethane dithiol
ELISA: enzyme-linked immunosorbent assay
EV: empty vector
FCS: fetal calf serum
FlAsH: fluorescine arsenical hairpin-binder
FLR: finger loop region
Gö6983: PKC inhibitor
GPCR: G protein coupled receptor
GRK: GPCR kinase
HA: haemagglutinin
HEK: human embrionic kidney cell
Iso: isoproterenol
KD: kinase dead
KO: knockout
M1R, M2R, M3R, M4R, M5R: human muscarinic 1,2,3,4, and 5 acetylcholine receptors
MOP: μ-opioid receptor
NanoLuc: Nano luciferase
ORF: open reading frame
PBS: phosphate buffered saline
PKC: protein kinase C
PMA: Phorbol 12-myristate 13-acetate
PTH(1-34): shortened parathyroid-hormone
PTH1R: parathyroid hormone 1 receptor
RFU: relative fluorescence units
SD: standard deviation
SDS: Sodium dodecyl sulfate
SEM: standard error of the mean
Squassh: segmentation and quantification of subcellular shape
V2R: vasopressin type 2 receptor
YFP: yellow fluorescent protein

